# Surface electrostatic networks control hydrophobic core remodeling in a pH-dependent switching protein

**DOI:** 10.64898/2026.07.16.738941

**Authors:** Iain M.H. McDonald, Luis B.P. Socas, Max Walton-Raaby, Xiaorong Liu, Sathish Dasari, Benjamin G. Emmanuel, Mahnoor S. Butt, Subha Kalyaanamoorthy, Charles L. Brooks, Elizabeth M. Meiering

## Abstract

Communication between protein surfaces and their buried cores is central to protein structure and function, yet this phenomenon remains challenging to predict and control at high resolution. Changes in the protonation of surface ionizable residues communicate with the hydrophobic core, for example, in diverse pH-dependent protein functions. Hisactophilin, a histidine-rich actin- and membrane-binding protein, provides a general model for exploring such communication as it exhibits a finely tuned pH-regulated myristoyl-switching function. Upon reversible proton binding, the myristoyl group shifts between being sequestered in the hydrophobic core and more solvent accessible. In the current study we utilize experimental and computational approaches we uncover how binding of ∼1.5 net protons alters electrostatic interactions involving ionizable residues distributed across much of the protein surface. These changes are transmitted to the hydrophobic core through dynamic communities of ionizable and hydrophobic residues which substantially rearrange upon switching. The effects of mutating individual ionizable residues are weaker than those of core hydrophobic residues, and only combined mutation of multiple ionizable residues caused substantial functional change. Together, these results reveal how communication between surface ionizable residues and the hydrophobic core is mediated by extensive interaction networks that reorganize in response to changes in protonation. These results may provide general insights for understanding protein cooperativity and the coupling of surface and core residues in protein function, disease, evolution, engineering, and design.

**Significance Statement:** How changes on the protein surface, such as proton binding to ionizable amino acids, are communicated to the protein core to regulate protein stability and function remains ill-defined. Synthesis of experimental and computational analyses resolves the distributed networks of surface ionizable residue interactions coupled to the hydrophobic core that control pH-dependent myristoyl switching in hisactophilin. Small changes in protonation that create and alleviate local electrostatic repulsion give rise to protein-wide changes in fluctuating surface-core interactions. This distributed electrostatics-core coupling mechanism may help explain the often underrecognized and long-range impacts of ionizable residues in proteins and provide a framework for interpreting the effects of mutations in fundamental and applied protein science.

## Introduction

Noncovalent interaction networks in proteins underpin their structure and function, yet they remain challenging to define and predict. Such interactions at the protein surface, and their coupling to the protein core, are of broad interest for understanding, predicting, and controlling allosteric regulation in nature, biotechnology, and medicine^1–10^. Allostery refers to processes whereby a perturbation, such as ligand binding or protonation, regulates a distinct functional site. Much has been learned about allostery, advancing from the lock-and-key model for switching between different protein states to an increasing recognition and definition of shifts in dynamic networks of interactions in different functional ensembles^8,9,11^. Owing to the intricacy of the communication pathways in proteins, however, a precise understanding and rational control of allostery is still a grand challenge^12,7,13^.

In the work described here we integrate experimental and computational methods to investigate how proton binding to surface ionizable groups drives myristoyl switching in hisactophilin, a histidine-rich actin- and membrane-binding protein, as a model system. Reversible proton binding regulates numerous biological processes through pH-dependent changes in protein structure, dynamics, and interactions^5,14–17^. Myristoylation is a common lipid modification whereby a C14 saturated fatty acyl chain is covalently linked to an N-terminal glycine in a protein^18,19^. Allosteric control of myristoyl switching plays a vital role in a wide variety of regulated protein-membrane and protein-protein interactions^19–21^. In hisactophilin, the myristoyl moiety is sequestered in the central hydrophobic core of the protein near neutral pH and becomes more solvent accessible at decreased pH during chemotactic signalling^22–25^. The binding of ∼1.5 protons with decreasing pH causes the protein to switch from the sequestered to accessible state^22^, making hisactophilin an intriguing model system to investigate how changes in surface electrostatics can govern the behaviour of the buried hydrophobic core (Fig. 1 a and b). The pH-dependence of switching is highly sensitive to mutations of hydrophobic residues, but the roles of ionizable groups in determining switching have not yet been well characterized. Solvent-exposed histidines in a surface β-turn, H89 and H91, were previously implicated in controlling switching via changes in protonation and electrostatic interactions with D57 and other nearby histidines^26^.

**Figure 1.**
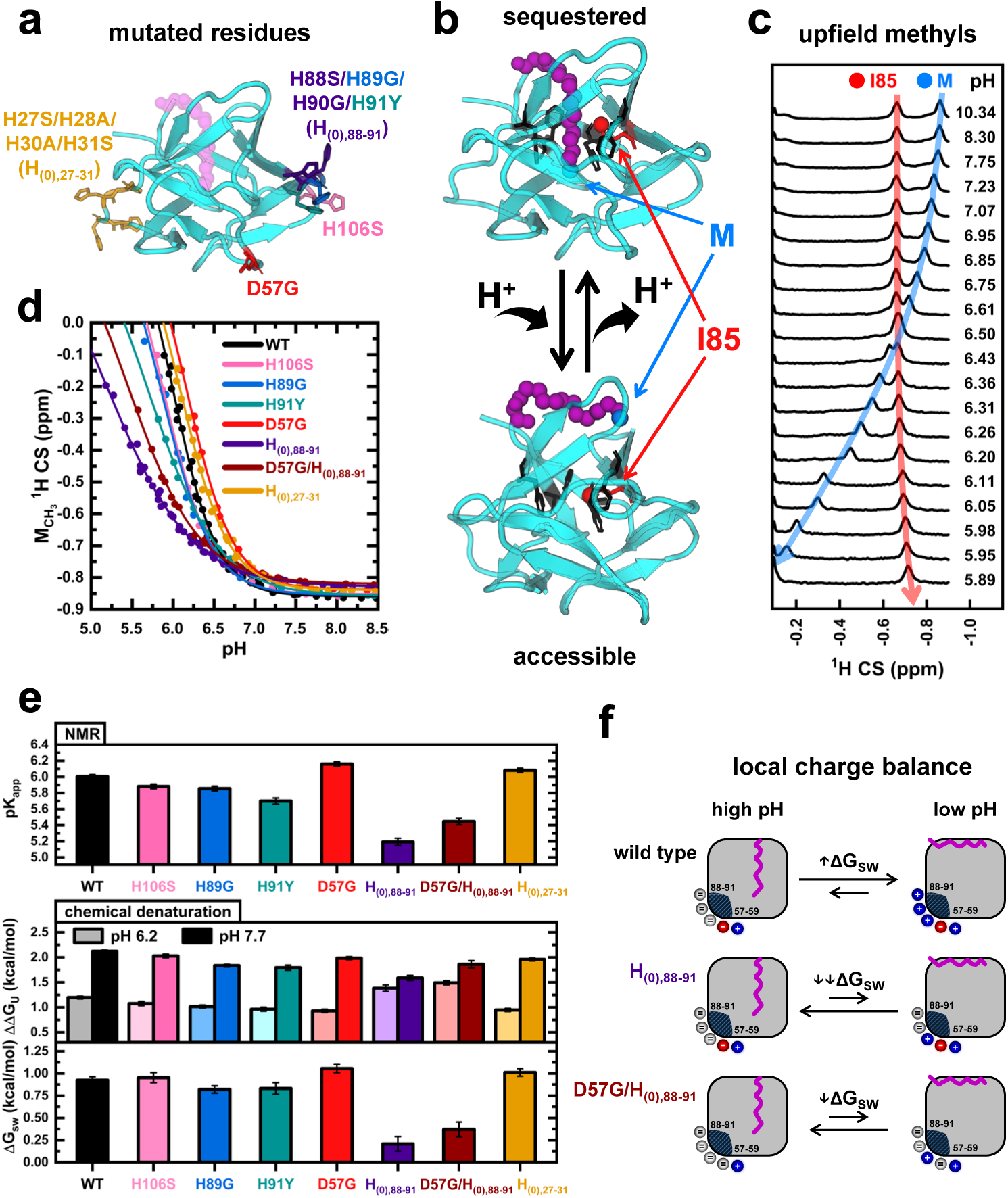
Experimental measurement of switching function in hisactophilin variants. (a) Location of mutated residues. (b) Illustration of the myristoyl group (purple) switching between sequestered and accessible states due to proton binding/release. Core aromatic residues are shown in black and I85 in red. The NMR trackable methyl groups for the myristoyl (M) and I85 are shown as blue and red spheres, respectively. (c) 1D-^1^H NMR spectra for wild-type hisactophilin showing the movement with pH of the methyl peaks due to switching. (d) pH-dependence of ^1^H chemical shift (CS) of the myristoyl methyl in hisactophilin variants. (e) Summary of the switching capability assessed by switching pK_app_ obtained by NMR (top bars) and by chemical denaturation-based thermodynamic cycle analysis (middle and bottom bars). (f) Schematic representation of the local charge balance between the H88–91 and 57–59 regions to explain mutant switching behavior.

To determine how surface ionizable groups regulate myristoyl switching, we integrated nuclear magnetic resonance (NMR) spectroscopy, mutational analysis, thermodynamic coupling measurements, constant-pH molecular dynamics (CpHMD) simulations and dynamic network analysis. We find that switching has low sensitivity to changing individual ionizable residues. Instead, the collective experimental results and state-of-the-art CpHMD analyses reveal that electrostatic allostery is distributed over many surface ionizable residues through state-specific fluctuating interaction networks that couple the protein surface to the hydrophobic core. Thus, our results uncover a general mechanism for electrostatic allostery mediated by dynamic networks that explain how numerous surface ionizable residues can collectively regulate the protein core.

## Results and Discussion

### Local charge balance is a major contributor to the pH-dependence of myristoyl switching

Identifying the ionizable residues responsible for pH-dependent myristoyl switching in hisactophilin is challenging because, as in many soluble globular proteins, these residues are numerous and distributed across the protein surface. Hisactophilin contains many ionizable residues, including 31 histidine, 9 lysine, 1 arginine, 6 aspartic acid, and 7 glutamic acid residues, all on the surface and comprising 46% of the primary sequence (selected ionizable groups shown in Fig. 1a). The protein undergoes a reversible conformational change coupled to pH, transitioning between a myristoyl-sequestered state near neutral pH and a myristoyl-accessible state in response to proton binding at decreased pH^22,26^ (Fig. 1b). This sequestered-accessible equilibrium can be sensitively monitored by NMR (Fig. 1c)^22^. Two isolated peaks observed in the 1D-^1^H NMR spectrum of hisactophilin at high pH correspond to the terminal methyl of the myristoyl group and the δ-methyl of I85, which are shifted upfield due to packing near aromatic residues in the protein core (Fig. 1b and c)^22^. With decreasing pH, the myristoyl methyl resonance shifts downfield as the acyl chain becomes more solvent-accessible. Because exchange between the sequestered and accessible states is fast on the NMR timescale, the observed chemical shifts report the population-weighted average of the states^22^, providing a quantitative measure of switching.

Residues D57, H89, and H91 were previously implicated as important contributors to switching^26^. In addition, H89 and H91, together with H88 and H90, form a β-turn between β-strands containing hydrophobic core residues I85 and I93, which strongly influence switching^26,27^. Here we tested to what extent switching depends on electrostatics in this region by making mutations that replace charged residues with neutral ones while minimizing alterations in stability (Fig. 1a and SI Methods). To examine progressively larger perturbations, we characterized single mutants D57G, H89G, H91Y, H106S (another histidine spatially close to H88–91), a quadruple β-turn mutant replacing all four β-turn histidines, H88S/H89G/H90G/H91Y (H_(0),88-91_), and quintuple mutant D57G plus H_(0),88-91_ (D57G/H_(0),88-91_). As an internal specificity control, we also examined a quadruple mutant replacing four histidines H27S/H28A/H30A/H31S (H_(0),27-31_) located on the opposite side of the protein.

To determine the effect of these mutations on switching, the pH-dependence of the myristoyl methyl chemical shift of the mutants is compared with wild type (Fig. 1d). A Hill equation fit of the pH titration curves yields values for the apparent pK (pK_app_), the pH at the midpoint of the switching process (Fig. 1e and Table S1). We also measured the switching process via a chemical denaturation-based thermodynamic cycle (SI Methods)^22,27,26^. This analysis uses the measured Gibbs free energy of unfolding (ΔG_U_) for myristoylated and non-myristoylated variants at high (7.7) and low (6.2) pH to obtain the free energy of switching ( ΔG_SW_ = ΔΔG_U,high pH_ − ΔΔG_U,low pH_), which reports on the extent of coupling between pH changes and myristoyl state. Decreased ΔG_SW_ indicates that this coupling is weakened by the mutation (Fig. 1e and Table S2).

The mutation series reveals that all variants retain switching capability but exhibit altered pK_app_ values. Point mutants H89G, H91Y, and H106S show a small shift in the titration curves to lower pH values, while replacement of the four β-turn histidines (H_(0),88-91_) markedly diminishes switching. Unexpectedly, D57G shifted switching to higher pH rather than lower pH. This mutation effect is also observed in the multi-mutant D57G/H_(0),88-91_, with the pH dependence again shifted to higher pH compared to H_(0),88-91_ (Fig. 1d and e). In contrast to the decreased pH-dependence of switching upon removing 4 histidines in H_(0),88-91_, removing 4 histidines in H_(0),27-31_ slightly increases it. Further, the effects of mutations on switching obtained by NMR are highly consistent with those measured by thermodynamic cycle analysis (Fig. 1e).

These results can be interpreted in the context of a net charge-based switching model (Fig. 1f). We hypothesized previously that switching in hisactophilin has two key features: 1) stabilization of the accessible state by formation of favourable electrostatic interactions at low pH (e.g. between D57 and H89/H91), associated with increased histidine pK_a_ values, and 2) stabilization of the sequestered state by decreased histidine pK_a_ values, lowering unfavourable repulsion among multiple spatially clustered positively charged residues in the vicinity of H89/H91^22,26^. Together, the mutant data in Fig. 1 indicate that the net charge on the protein surface near H88–91 is of greater functional importance than the formation of specific pair-wise electrostatic interactions between the mutated residues. This finding is in line with growing recognition that allostery often goes beyond well-defined, short pathways and is better described by changes in more distributed protein interactions^7,9,28^. Such a mechanism is supported by other NMR and mutant results for hisactophilin that provide evidence for much of the protein participating in switching^26^. This more comprehensive distributed electrostatic hypothesis is also supported by the consistent relationship between local charge and switching pK_app_: increasing positive charge (as in D57G relative to wild type, and D57G/H_(0),88-91_ relative to H_(0),88-91_) raises switching pK_app_ whereas decreasing the positive charge in this region has the opposite effect (as in H89G, H91Y, H106S, H_(0),88-91_). Notably, the four histidines in H88–91 are surrounded by numerous other ionizable groups, such as those in the 57–59 turn (D57, H58, and K59), as well as H106 (Fig. 1a). In this distributed electrostatics framework, repulsion between spatially grouped positive residues in the sequestered state with decreasing pH drives switching via rearrangement of many surface electrostatic as well as core interactions (developed further below). The repulsive interactions go beyond those involving just H88–91, evidenced by residual switching in H_(0),88-91_. The removal of a similar set of 4 histidines in the control mutant H_(0),27-31_ has little effect on switching, indicating that the effects of both H_(0),88-91_ and D57G/H_(0),88-91_ are also related to their structural context in the protein and are not merely the result of removing four clustered histidines.

### Regions beyond H88–91 contributing to switching: re-wired networks in H_(0),88-91_

Although mutation of the H88–91 β-turn substantially diminishes the pH-dependence of switching, the H_(0),88-91_ mutant retains considerable switching competence. To further investigate the transition at residue level resolution, we measured pH-dependent backbone chemical shift perturbations (CSPs) by ^15^N–^1^H Heteronuclear Single Quantum Coherence (HSQC) NMR for wild type and the H_(0),88-91_ mutant (Fig. 2a, S1, and S2). At a given pH, wild type shows larger and more localized perturbations than the mutant, particularly near residue 56 and the 88–91 β-turn. However, under conditions that produce equivalent degrees of switching, as measured by the myristoyl methyl chemical shift (Fig. 1d), CSP magnitudes in H_(0),88-91_ approach those of wild type. These results indicate that local changes in residue environment are strongly related to the sequestered/accessible equilibrium rather than pH alone. Interestingly, in the absence of the β-turn histidines, enhanced perturbations appear in alternative regions, particularly around residues 68–73 (Fig. 2a), suggesting that the mutant reaches similar switching end states through altered surface interactions. Nearby ionizable groups, including D57, H58, K59, H68, and H71, may therefore adopt a more significant role in sensing pH changes in the absence of histidines at residues 88–91 (see below).

**Figure 2.**
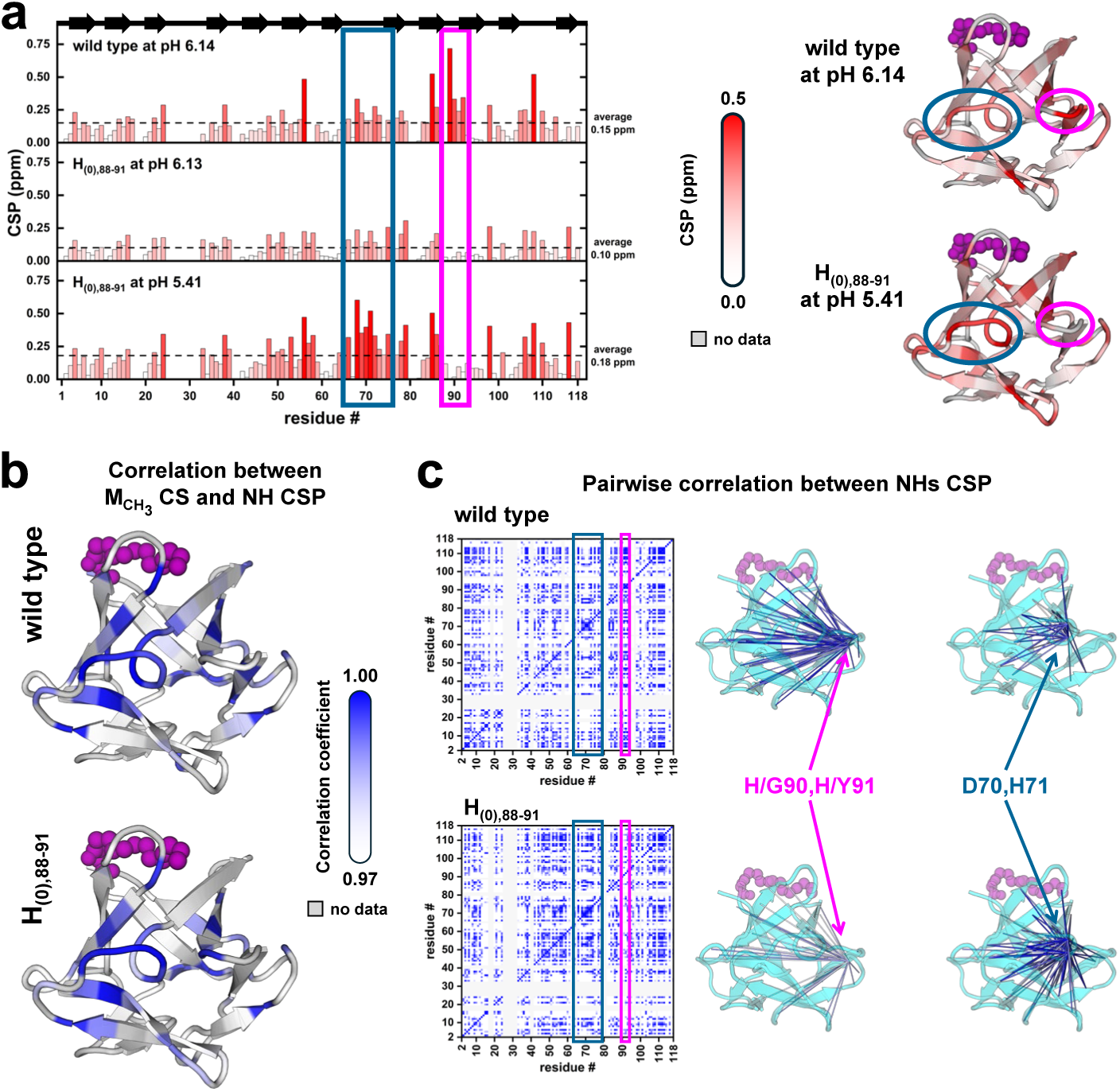
Chemical shift perturbation (CSP) analysis of wild type and the H_(0),88-91_ mutant. CSP is calculated using combined ^15^N and ^1^H chemical shifts as CSP = ([Δδ_H_^2^ + (Δδ_N_/5)^2^]/2)^1/2^, where Δδ is the perturbation relative to pH 8. (a) CSP values of wild type and H_(0),88-91_. pH 6.14 and pH 5.41 correspond to similar myristoyl methyl chemical shifts for wild type and mutant (Fig. 1d). The 70s loop region and 90s loop region are highlighted with blue and pink box and oval, respectively. (b) Pearson correlation coefficient between NH CSPs at each pH relative to pH 8 and the chemical shift of myristoyl CH_3_. (c) Pairwise correlation coefficient between each residue NH for both wild type and H_(0),88-91_. Lines in the ribbons show intramolecular correlations for residues 90–91 and 70–71. Intensity of colour for correlations increases with increasing R (see SI Methods)^28^.

To gain further mechanistic insight into switching, we correlated residue-specific amide CSPs with the myristoyl methyl chemical shift over the studied pH range. This analysis provides a measure of how strongly each residue is coupled to the global conformational transition. For both hisactophilin variants, most residues exhibit strong correlations with the global switching coordinate (Fig. 2b, S3a and b), indicating that the local perturbations reflect redistribution of the global conformational equilibrium not just independent local structural changes. The fact that most correlations remain similar in wild type and H_(0),88-91_ is consistent with the underlying transition being preserved upon mutation. In addition, we analyzed pairwise NH chemical shift and CSP correlations, which is useful for sensitively mapping communication pathways underlying allosteric transitions^29,28^. We observed a clear redistribution of correlated responses in the H_(0),88-91_ mutant relative to wild type (Fig. 2c and S3c), suggesting re-wired patterns of residue-residue coupling during the pH-dependent transition. Specifically, residues 90–91 show less correlation with the rest of the protein in H_(0),88-91_ than in wild type, consistent with the removal of the β-turn histidines weakening their integration into the switching process. In contrast, residues D70 and H71 show higher correlations with other regions of the protein, suggesting potentially increased participation in switching. These changes demonstrate that H88–91 are not solely responsible for controlling switching and that the coupled residue network is sufficiently plastic to reorganize in response to mutation (Fig. S3c). These results provide evidence that the surface interactions involved in driving switching are adaptable: when the β-turn histidines are removed, neighboring loops become more actively engaged in mediating the conformational rearrangements required for switching. Together with the generally limited effects of point mutations on switching energetics, these findings support the involvement of extensive electrostatic networks contributing to allostery, which we examine further below.

### Surface protonation is accompanied by hydrophobic core contraction

To gain deeper insight into how protonation on the surface of the protein is connected to the core, we further examined 1D-^1^H NMR titration data. In particular, the ring currents of core aromatic residues also affect the hydrophobic residue I85^22^, which, unlike the myristoyl group, remains buried in both the sequestered and accessible states. Interestingly, the I85 δ-methyl resonance shifts further upfield with decreasing pH (Fig. 1b), consistent with increased aromatic shielding. This observation suggests that protonation induces tighter packing of the hydrophobic core in the accessible state. Notably, a similar magnitude upfield shift is also observed in non-myristoylated hisactophilin (Fig. 3a), consistent with core contraction being related to surface protonation and not solely a consequence of myristoyl displacement. It is noteworthy that this upfield movement of I85 δ-methyl chemical shift is less prominent for the H_(0),88-91_ and D57G/H_(0),88-91_ multi-mutants than for wild type, but in the absence of the myristoyl group their pH dependence becomes very similar, with pK_app_ values ∼6.8 (Table S3). This can be understood in terms of an intrinsic response of the core to contract due to protonation on the surface, which occurs more easily in the non-myristoylated variants. However, when the myristoyl group is present, the pK_app_ of this core response is shifted, indicating that the buried acyl chain resists core contraction and thereby effectively lowers the pK_app_ of the process (Fig. 3a schema).

**Figure 3.**
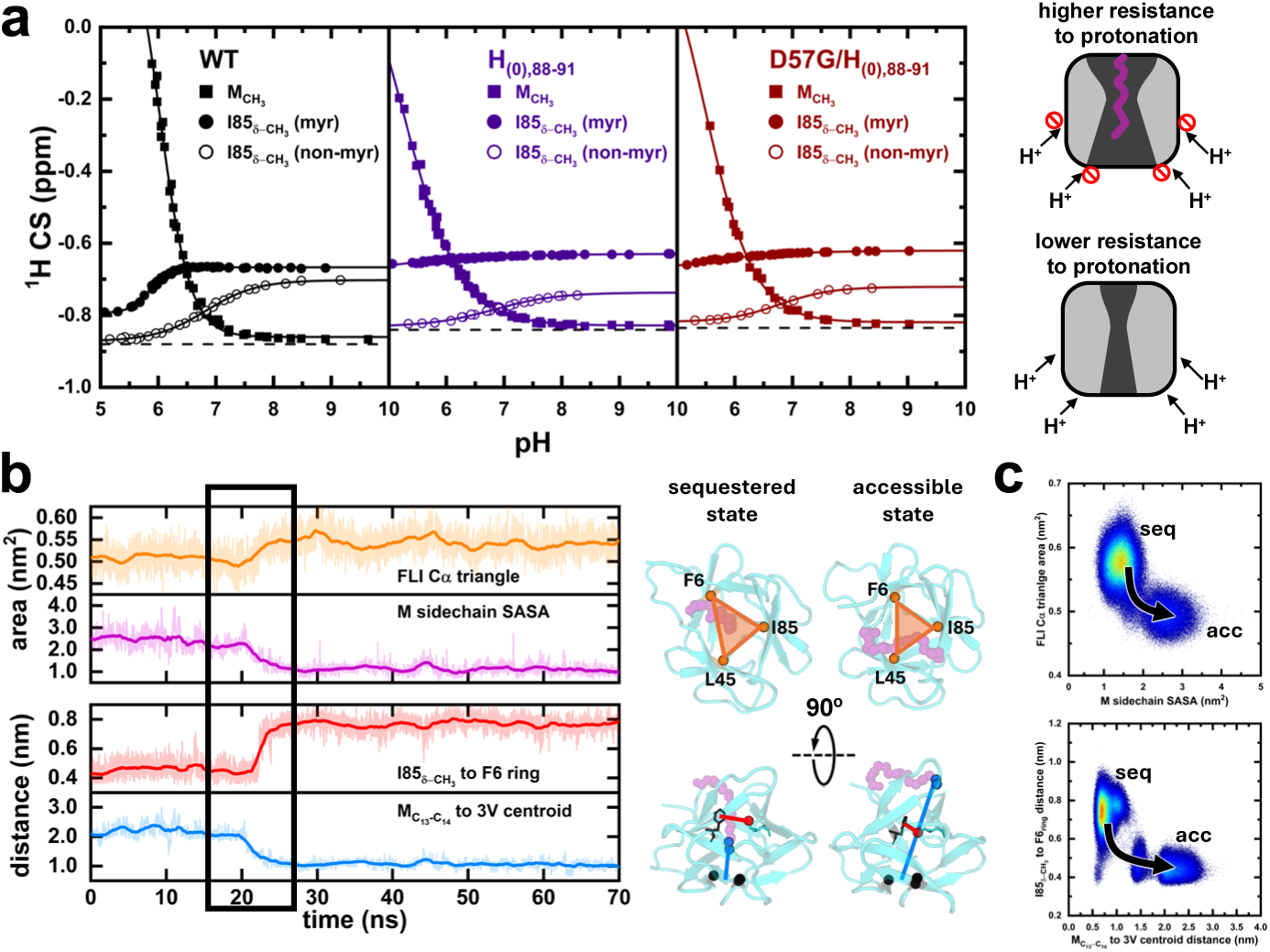
Structural changes in hisactophilin core due to surface protonation. (a) Experimental ^1^H chemical shift (CS) of myristoyl CH_3_ (squares) and the δ-CH_3_ of I85 sidechain (circles) for myristoylated (filled symbols) and non-myristoylated (open symbols) variants of wild type, H_(0),88-91_, and D57G/H_(0),88-91_ mutants. The schema (right) summarizes coupling between surface protonation and core rearrangement. Red dashed circles indicate decreased protonation when the myristoyl (purple) is sequestered in the core. (b) Representative time evolution of several structure metrics during constant pH molecular dynamic simulations. Metrics are summarized in the ribbons representations, namely: (top) solvent accessible surface area (SASA) of the myristoyl acyl chain (purple); and the area of the triangle formed by the Cα of F6–L45–I85 (orange); the conformational change during the simulation is depicted with a rectangle. (bottom) the distance (blue line) between the end of the myristoyl group (midpoint of C_13_-C_14_ in blue spheres) to the centroid of the Cαs of V21–V61–V101 (3V, black spheres); the distance (red line) between I85 δ–CH_3_ (red sphere) and the center of the F6 ring (black sticks). The conformational change during the simulation is depicted with a rectangle. (c) Plots of the measured structural parameters using data from all 160 simulation replicates. Arrows indicates the conformational changes from the sequestered (seq) to accessible (acc) states.

To examine this mechanism at atomic resolution, we performed λ-dynamics CpHMD simulations in explicit solvent for myristoylated hisactophilin^14,30–34^. Unlike conventional MD simulations, λ-dynamics CpHMD treats protonation as a dynamic variable, enabling accurate modeling of coupled protonation and conformational equilibria that govern pH-dependent switching. Two starting structures were used in the simulations, representing the sequestered and accessible conformations, and simulations were conducted at high (7.7) and low (6.2) pH (see SI Methods). Fig. 3b shows a representative MD trajectory including a transition between accessible/sequestered conformational states. To assess the myristoyl position, we measured the distance from its C_13_–C_14_ bond to the centroid of Cα atoms of valines 21, 61, and 101 (3V, Fig. 3b blue curve)^26,35^, as well as the myristoyl chain’s solvent accessible surface area (SASA, Fig. 3b purple curve). In addition, Fig. 3b shows the distance between the δ-methyl of I85 and the center of the F6 ring, located on opposite sides of the core (red curve), and the area of the triangle formed by the Cα of three core residues F6, L45, and I85 (FLI triangle, orange curve). These calculations illustrate that transitions between states are accompanied by concerted changes in all these parameters.

To ensure the observations in Fig. 3b are robust, we performed a combined analysis of the structural data from 40 replicate simulations (100 ns per replicate) for each starting state (Fig. 3c and S4). The accessible state consistently exhibits reduced I85–F6 distance and decreased FLI triangle area, which agrees with the core being contracted, providing structural explanation for the NMR observations. The core contraction upon switching is an interesting observation in the context of the hisactophilin’s biological function. Experimental data show that the overall structure of the protein is maintained when the myristoyl becomes exposed for membrane binding^24–26^. Thus, closer packing of hydrophobic core residues upon myristoyl exposure may be important for maintaining the structure required for both membrane and actin binding. Measuring the distance between the end of the myristoyl group and the 3V centroid showed a low-populated state (∼9% of the frames) between the two end states (with intermediate M_C13-C14_ to 3V centroid distance of ∼1.4 nm, between ∼0.7 nm and ∼2.1 nm for sequestered and accessible, respectively (Fig. 3c and S4). This is in line with previous simulations, which also reported a partially accessible state^26^. For the scope of this work, we focus on the differences between the end states of switching; therefore, all subsequent CpHMD analyses consider frames in which the M_C13-C14_–3V distance was classified as either accessible or sequestered (see SI Methods).

### Switching is driven by differences in state-dependent surface dynamics and surface-core coupling

#### Surface dynamics (RMSF) and local coupling

We next explored how surface electrostatics are linked to conformational switching in hisactophilin, with a focus on the dynamic role of histidine residues, which have been implicated in ionizing over our working pH range^22^. For comparisons between the sequestered and accessible states we used pH 6.2, where both conformations are significantly populated (Fig. 1); generally similar but attenuated effects are observed at pH 7.7. The structural parameters that distinguish the accessible and sequestered states (Fig. 3) are accompanied by changes in the protonation profile (Fig. 4b and S5). The protonation predicted by CpHMD (Fig. 4b) corresponds closely to the experimental values in average net value (Fig. 4a) and in the small differences upon switching (∼1.5 proton increase from sequestered to accessible)^22^. The net protonation is shifted and decreased in H_(0),88-91_ (Fig. S6), consistent with results for this variant in Fig. 1–3. This demonstrates remarkable accuracy can be obtained by simulations even for proteins with large numbers of ionizable residues, supporting more detailed analyses of simulation results described below. In addition, the degree of state-dependent individual protonation differences is extensive, despite the small net change, in line with many histidines participating in the switching process (Fig. S5).

**Figure 4.**
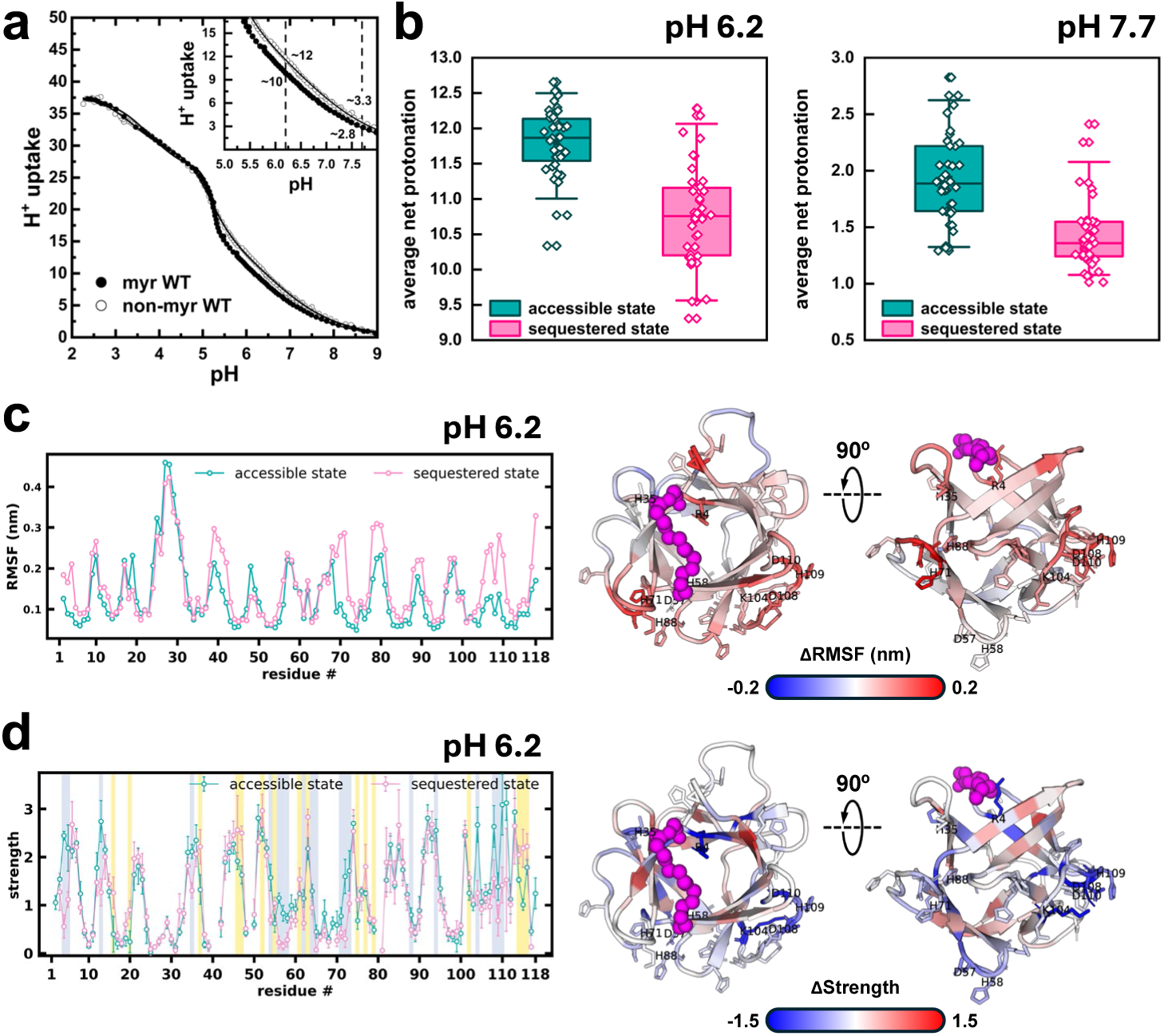
Changes in protonation and dynamics of hisactophilin due to switching. (a) Hisactophilin proton uptake measured experimentally by HCl titration of myristoylated (filled symbols) and non-myristoylated (open symbols) variants. (b) Box-and-whiskers plots of the net protonation of histidine residues obtained by CpHMD at pH 6.2 and 7.7 for the sequestered and accessible states simulations (see SI Methods for details). Diamonds correspond to the mean protonation of each replicate; the box signals the mean and quartiles while the whiskers correspond to the 5 – 95% range. (c) RMSF values for accessible and sequestered state simulations at pH 6.2, alongside ribbon representations of the structure depicting the ΔRMSF (sequestered – accessible). (d) Strength values (the sum of a given node’s |DCC| edge weights) for the same simulations, alongside ribbon representations of the structure depicting the ΔStrength (sequestered – accessible). Residues with statistically significant (q<0.10) differences are highlighted in yellow or grey for values larger in the sequestered or accessible states, respectively. Residues shown as sticks in both (c) and (d) ribbons correspond to those with differences in both RMSF and Strength. Positive ΔRMSF (red in c) tends to be inversely correlated with ΔStrength (blue in d) (Fig. S9).

Beyond protonation differences, the two states also exhibit pronounced differences in structural flexibility. Root-mean square fluctuation (RMSF, see SI Methods) identifies five predominantly surface-exposed regions (residues 38–44, 68–74, 77–83, 86–92, and 104–112) that undergo marked reductions in flexibility upon transitioning from the sequestered to the accessible state (Fig. 4c and S7). Interestingly, these regions form a largely contiguous surface spanning a large portion of the protein structure, with the 88–91 β-turn region serving as a bridge between other segments of structure, consistent with large scale coordination rather than localized fluctuations.

To further investigate this coordination, we generated interaction networks of our systems. Residues and the myristoyl are represented as nodes, while interacting nodes are connected by edges weighted by their pairwise absolute dynamic cross-correlation (DCC) values (Fig. S8, see SI Methods for details). Using Strength (the sum of a node’s |DCC| edges) to quantify the extent of local dynamic coupling, we found that Strength and RMSF present an overall inverse relationship (Fig. 4d and S9), indicating that stronger dynamic coupling contributes to reduced flexibility. Notably, we observe several ionizable residues exhibit significantly altered Strength values between states and their Strength primarily increases upon transitioning from the sequestered to the accessible state (R4, H35, D57, H58, H65, H71, H88, K104, D108, H109, and D110, among the grey highlights in Fig. 4d, Tables S5 and S6). Moreover, summing the Strength of electrostatic residues with significant state-dependent differences increased from 20.8 in the sequestered state to 26.1 in the accessible state, further supporting the enhanced dynamic coupling of surface electrostatics in the accessible state.

#### Community membership and flux

Both experiment and simulation strongly suggest that switching cannot be attributed to a select few residues in hisactophilin. Instead, the global and cooperative nature of the process requires a holistic description that accounts for the collective influence of all residue interactions. Toward this goal, we applied the Leiden community detection algorithm^36^ to the residue interaction networks to identify 1) communities in each conformational state, and 2) how these communities reorganize upon switching. We discover markedly different consensus community maps in the sequestered and accessible states at pH 6.2, uncovering the coupling between ionizable surface residues, their coupling to the core, and the interactions that drive switching (Figs 5a–h, S10, and Table S4).

**Figure 5:**
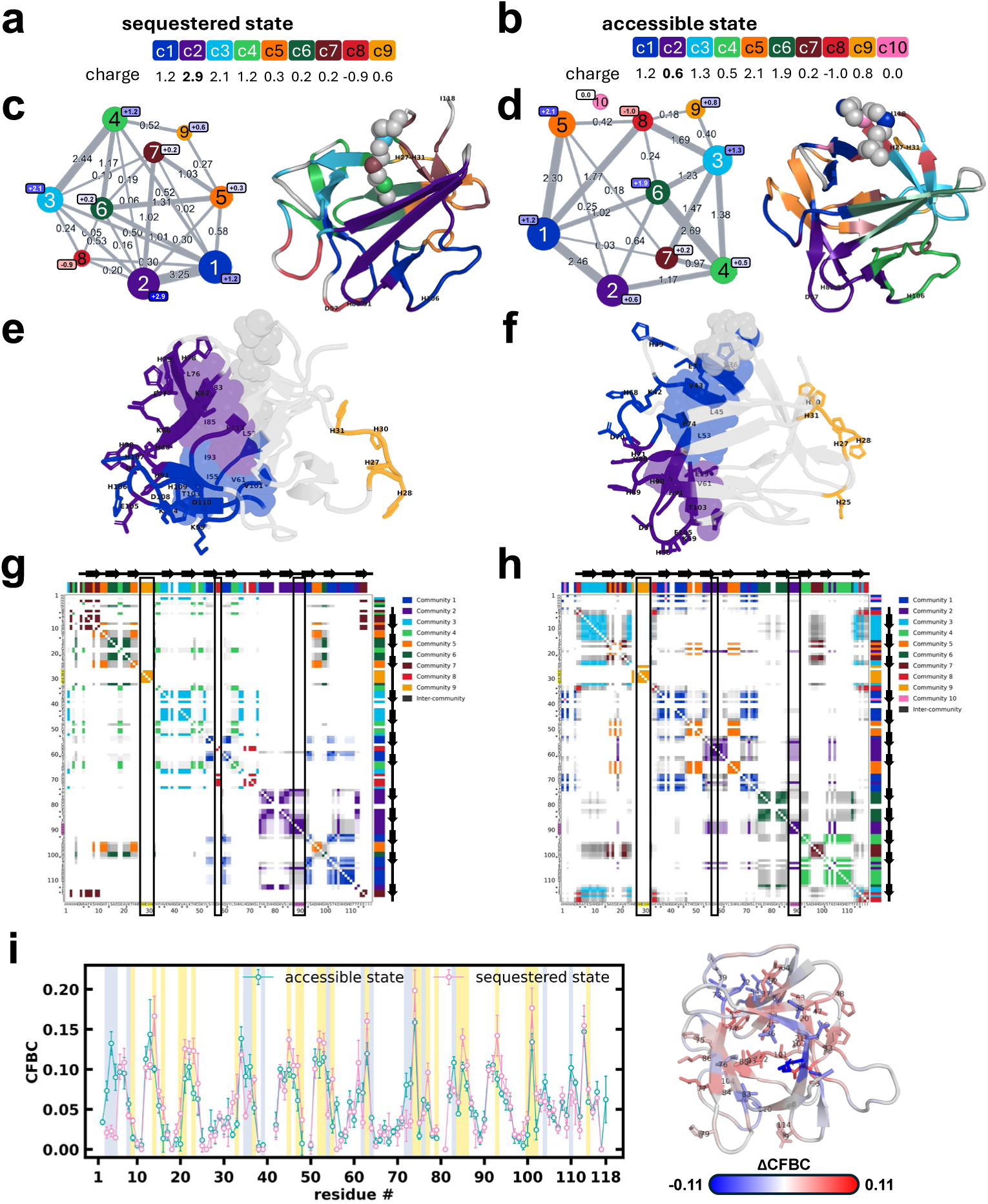
Community analysis of hisactophilin simulation trajectories at pH 6.2. (a) and (b) Communities (residues listed in Table S4) are numbered according to their size from c1, the community with the largest number of residues (nodes), to c9/10, with the smallest number of nodes. The net charge of each community is given (see SI Methods for details), with the value for c2 (containing H88*–*91) in bold. Note: I118 is not assigned to a community in the sequestered state. (c) and (d) Community-based graphs showing intercommunity communication for sequestered and accessible states, respectively. Circle sizes are proportional to community sizes and edge widths are proportional to the sum of edge weights between communities. Community net charge is given in the squares adjacent to the nodes, coloured blue and red for positive and negative charge, respectively, with intensity of colour increasing with charge magnitude. Adjacent coloured ribbon representations show communities in the protein structure. (e) and (f) Structural representation of communities 1 and 2 for sequestered and accessible states, respectively. Surface ionizable residues are shown in sticks and core residues in space-fill. (g) and (h) Pairwise node co-occupancy consensus matrices for sequestered and accessible states, respectively. Communities are coloured as in panels a-d, while gray denotes node pairs with intercommunity crossover, with darker shading for higher co-occupancy. Rectangles on the matrices highlight regions of mutated residues: 27–31, 57, 88–91, which show little (27–31) or extensive (57, 88*–*91) changes between states. (i) Current-flow betweenness centrality (CFBC) values for the sequestered and accessible states (Table S5). Significant differences are highlighted on the plot in yellow (grey) for residues where CFBC value is significantly larger (q<0.10) in the sequestered (accessible) state than in the accessible (sequestered) state (Table S6). Ribbons representation of the difference in CFBC between sequestered and accessible states, ΔCFBC (sequestered – accessible), is coloured from red (higher in sequestered) to blue (higher in accessible); residues with significant differences are also shown as sticks labeled by residue number.

To investigate the collective nature of hisactophilin’s interactions, we quantified the extent of intercommunity communication (Fig. 5c and d) and net community charge, contextualizing whether interactions are predominantly attractive or repulsive (see SI Methods). The sequestered state features enhanced communication between highly positive net charge communities. Most notably in the sequestered state, communities 1 and 2 (containing H106 and H88–91, respectively, which diminish switching when mutated, Fig. 1) share the strongest edge among all community pairs. Conversely, while comprised of different residues in each state, community 8, the only negatively charged community, forms markedly stronger edges with positive net charge communities in the accessible state (summed edge weights are 4.30 for accessible and 0.79 for sequestered). These favourable interactions may abate the overall repulsion from the high net positive protein charge at pH 6.2 (Fig. 4) and facilitate increased protonation, consistent with the experimentally observed increase in pK_app_ values for the accessible state^22^.

Experimental measurements identify H88–91 as playing a major role in mediating pH changes. In line with this observation, community 2 (containing H88–91) undergoes substantial reorganization between the sequestered and accessible states, exhibiting a distinctly higher net charge in the former (+2.9 versus +0.6, Fig. 5a and b). A similar trend is observed for the community containing H106 (sequestered community 1 +1.2 versus accessible community 4 +0.5), also experimentally linked to switching (Fig. 1). In contrast, community 9 containing H27, H28, H30, and H31 shows minimal change in its constituents, net charge, and intercommunity communication between states, directly mirroring our experimental results demonstrating its functional decoupling from the switching mechanism. The accessible state features a more balanced community charge distribution and weaker edges between communities with substantial positive net charge. Conversely, apparent electrostatic frustration in the sequestered state likely favours the conformational change to the accessible state, redistributing charges into a more energetically favourable network.

Moreover, the community analysis provides crucial insights into the intricacies of dynamic surface–core coupling, as many of the communities include both ionizable surface and hydrophobic residues (Fig. 5a–h and Table S4). For example, community 2 in the sequestered state contains both H88–91 and five core residues including L76 and I85. We show that repacking of I85 upon surface protonation is a feature of switching (Fig. 1 and 3), and previous experiments demonstrated that mutations of I85 and L76 strongly impact switch energetics^26^. Likewise, the sequestered H106 community (1) also contains I93, which also modulates switching^26^. Many prominent community rearrangements between states involve ionizable residues mutated here, or previously mutated hydrophobic residues, that impact switching. Interestingly, about two-thirds of the residues exhibiting elevated RMSF in the sequestered state (Fig. 4c) belong to communities 1 and 2. In contrast, H27–H31 show slightly decreased RMSF and participate in a small community that undergoes little change in membership and lacks hydrophobic residues, consistent with its minimal contribution to switching.

#### Surface-core communication pathways

Further, to identify the nodes mediating network-wide communication and how they differ between states, we calculated current-flow betweenness centrality (CFBC, see SI Methods)^37^. We uncover a global re-routing of communication dictated by the myristoyl environment (Fig. 5i), aligning with the hypothesis that switching is governed by a delicate balance of numerous stabilizing and destabilizing interactions^26^. Intriguingly, the residues with the largest CFBC values in both states also tend to display low flexibility (Fig. S11), consistent with rigid structural elements serving as effective transmitters for long-range communication^38,39^.

Across all systems, we observe much larger CFBC values for core and hydrophobic residues than for surface electrostatic residues (Fig. 5i and Table S5). Comparing between the myristoyl states at pH 6.2, the residues in the upper portion of the myristoyl pocket (R4, V36, L76, and V83) show increased CFBC values and decreased RMSF in the accessible state, likely due to closure of the cavity and closer packing (consistent with Fig. 3), providing an efficient route of communication across the protein and to the myristoyl. Conversely, the sequestered state yields larger CFBC values for deep core residues lining the lower myristoyl pocket (L14, L45, F74, I85, I93, and V101). This, in addition to the modest CFBC for the myristoyl nodes, suggests that communication may primarily travel around the myristoyl group, rather than through it, positioning the myristoyl as a mediator of strain (leading to enhanced RMSF) rather than a communication hub. This view is reinforced by the incipient switching of non-myristoylated hisactophilin observed experimentally (Fig. 3a). Among other residues in the sequestered state at low pH, I85 and I93 are critical communication conduits and known modulators of myristoyl switching^26,27^. Moreover, these two residues are linked by the frustrated H88–91 β-turn, which our work proves to be important in switching. The surface electrostatic residues also tend to increase in CFBC in the sequestered state, which together with this state’s low Strength for these residues and higher proportion of repulsive intercommunity communication implies a more constrained surface electrostatic network with fewer viable pathways. Overall, the myristoyl sequestration substantially reroutes communication relative to the accessible state: deep core residues become essential for transmitting the protonation signals received by structurally coupled surface community members to neighboring residues throughout the myristoyl binding pocket, providing the key link between surface and core in driving pH-dependent switching.

### Molecular mechanism of pH-dependent switching in hisactophilin

Based on synthesis of experimental and computational studies, we propose that distributed electrostatic allostery, featuring electrostatic repulsion, controls pH-dependent myristoyl switching in hisactophilin (Fig. 6). Switching arises from dynamic communities, comprised of ionizable and hydrophobic residues, which encompass much of the protein and rearrange extensively between the myristoyl sequestered and accessible states to rebalance electrostatic networks and hydrophobic packing (Fig. 5). In the sequestered state, proton binding is disfavoured by an increased localization of positively charged surface residues in two large communities, constituting a wide swathe around four histidines (88–91) in a β-turn. Electrostatic repulsion is propagated through networks of ionizable residue and into the core, manifested by increased fluctuations (RMSF) across the bulk of the protein (Fig. 4). This strain is abated in the accessible state by dispersing repulsive interactions, increased attractive electrostatic interactions (Fig. 5), and contraction of hydrophobic core residues capped by the myristoyl (Fig. 3).

**Figure 6:**
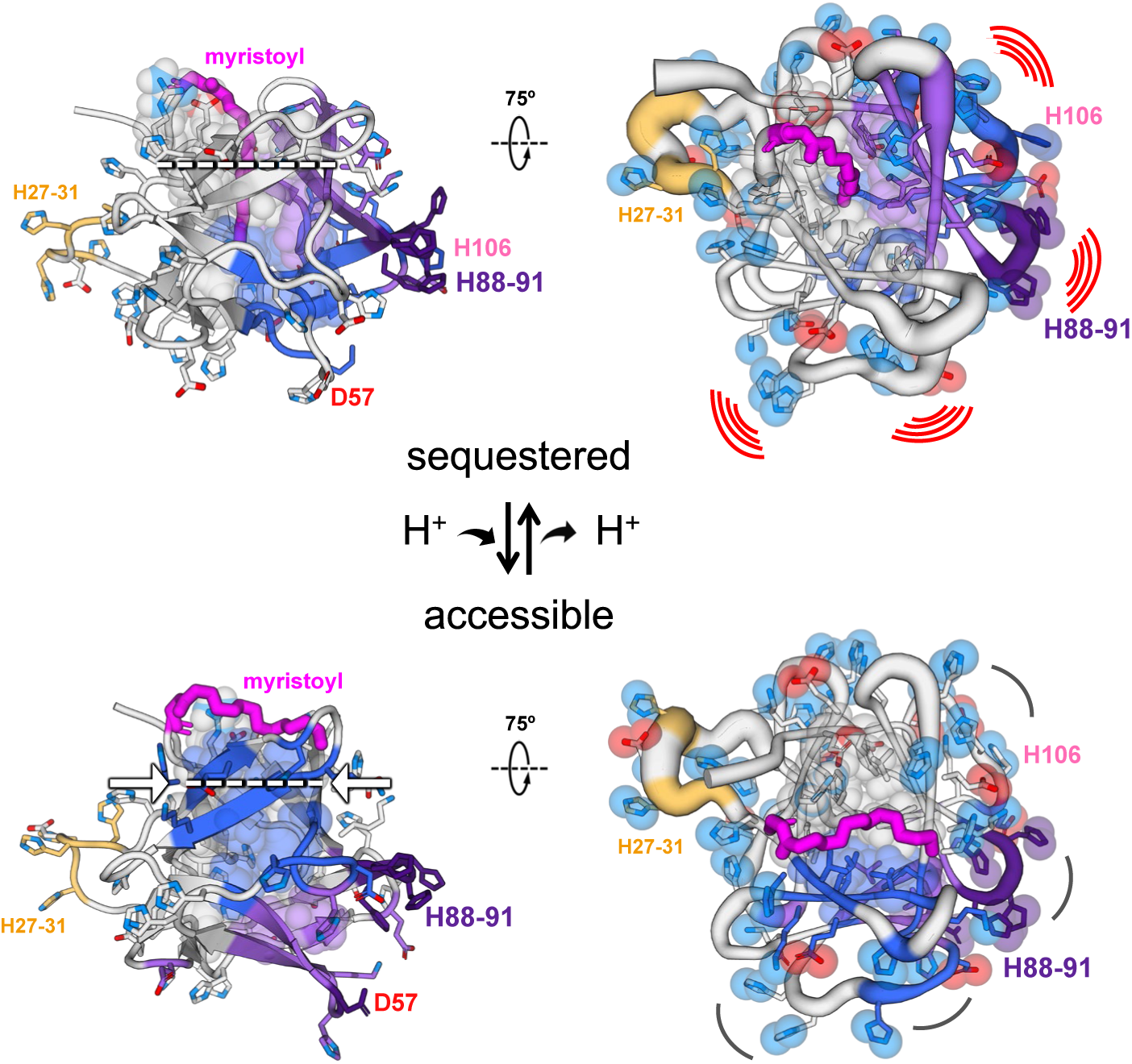
Mechanism of pH-dependent switching in hisactophilin by distributed dynamic community-based surface electrostatics coupling to hydrophobic core. Summary of features driving switching between the sequestered state (top) and the accessible state (bottom). Myristoyl, mutated residue labels, and the largest communities are coloured (1 in dark blue, 2 in purple, mutated residues in a deeper shade) as in preceding figures. Backbone shown in tube representation (right panels) with width proportional to RMSF (Fig. 4c). Arcs (right panels) indicate electrostatic repulsion and enhanced communication between communities 1 and 2, and 3 and 4, in sequestered. Ionizable residues are shown in stick representation, with spheres (right, for N and O atoms, coloured for non-mutated residues in lighter blue and red, respectively). Hydrophobic core residues are shown in space-filled representation. **Mechanism:** the sequestered state (top) has an expanded core to accommodate the myristoyl group. Multiple communities, including those containing H88–91 and H106, have increased RMSF compared to the accessible state (right panels). Increased motion of the surface is felt by structurally coupled core residues lining the myristoyl binding pocket, leading to a concerted change of the surface and core that minimizes electrostatic repulsion while simultaneously contracting the core (left, dashed white line and arrows). The decreased repulsion in the accessible state (bottom) is associated with increased proton uptake and reduced RMSF, as well as extensive rearrangements of communities (see also Fig. 5). Thus, switching is controlled by protein-wide coupled fluctuating electrostatic-hydrophobic interactions.

This mechanism is supported by multiple lines of evidence. 1) Mutations of ionizable residues in the vicinity of H88–91 decrease the pH-dependence of switching when net charge is decreased and increase when net charge is increased (Fig. 1 and 3a). 2) Analysis of chemical shift perturbations for amides throughout wild type and H_(0),88-91_ indicate electrostatic interactions are plastic and continue to mediate substantial switching even when these four histidines are absent (Fig. 2). 3) CpHMD simulations identify concerted structural changes throughout the protein upon switching (Fig. 3b and c) are associated with small changes in net charge that closely match experiment and involve many histidines (Fig. 4a, b, and S4). 4) Dynamic network analysis provides a detailed view of protein communication that is consistent with the experimentally demonstrated key roles of residues in the vicinity of H88–91 and minimal involvement of H27–31 in core coupling and switching (Fig. 1 and 5). Together these finding show that the binding of only ∼1.5 net protons gives rise to reorganization of protein-wide interactions.

These findings for hisactophilin align with previous observations and provide broader mechanistic insights into the roles of distributed surface electrostatic interactions in proteins. Surface electrostatic interactions have likewise been found individually to have smaller effects on proteins than core interactions^40,41^. Nevertheless, their individual and cumulative effects can be substantial and propagate widely throughout proteins to shape stability and allostery^5,42–47,6^. The significant effects of ionizable residues have often been underrecognized due to their small magnitude, transience, and relatively longer range. By analogy with our results, ionizable residues in distributed networks may help explain the prevalent and commonly modest impacts of surface, compared with core, substitutions on protein stability, function, and pathogenicity, such as those identified by surface and deep mutational scanning^48,1,3,49,50^. These deep mutational scanning experiments have recently discovered many previously unknown allosteric effects for surface ionizable residues in a variety of proteins^1,3,49,50^, such as KRAS which contains multiple surface allosteric pockets over a large proportion of the protein surface^47^. Understanding such allosteric sites is important for interpreting the effects of mutations in evolution and disease, and for developing next-generation allosteric therapeutics. Because these interactions are distributed and dynamic, their description requires modern computational simulations and analyses to capture correlated motions, communication pathways, and transient interaction communities to elucidate mechanisms^51,52,11,5,33,34^. By applying these approaches, state-of-the-art representations of ionizable residues can now yield specific insights with high accuracy and resolution. Taken together, these findings suggest that identifying distributed networks of surface ionizable residues, as demonstrated here for hisactophilin, may advance the understanding, prediction, and engineering of electrostatics in many other proteins.

## Materials and Methods

Brief descriptions of methods are below with additional details given in the SI Appendix.

### Mutant design and protein preparation

Surface ionizable residues were replaced with nonionizable residues based on sequence and structural information and protein-stability predictions. Wild-type and mutant hisactophilin variants were prepared as described previously^26^.Mutations and myristoyl incorporation were verified by mass spectrometry, and preservation of the global protein fold was evaluated by NMR spectroscopy.

### Protein stability and pH-dependent switching

Protein stability was measured by urea-induced denaturation monitored using intrinsic tyrosine fluorescence at pH 6.2 and 7.7. Denaturation curves were analyzed using a binomial extrapolation model, and the free energy of myristoyl switching was determined from the differences in unfolding free energy between myristoylated and non-myristoylated proteins using a thermodynamic-cycle approach^27^.

pH-dependent myristoyl switching was monitored by one-dimensional ^1^H NMR spectroscopy using the chemical shifts of the myristoyl terminal methyl group as probe of the accessible and sequestered states. Residue-specific responses were examined using ^1^H–^15^N HSQC titrations of isotopically labeled proteins. Amide chemical shift perturbations and pairwise chemical shift correlations were used to identify residues displaying coordinated pH-dependent responses. Proton uptake by myristoylated and non-myristoylated proteins was measured by potentiometric titration relative to protein-free controls.

### Molecular dynamics simulations

Initial conventional MD simulations were performed at pH 6.2 and 7.7 to obtain representative accessible and sequestered conformations of myristoylated hisactophilin. These structures were subsequently examined using CpHMD simulations, in which all 31 histidine residues dynamically sampled their protonation states. Four conformation–pH scenarios were simulated, corresponding to accessible and sequestered structures, each at pH 6.2 and 7.7. For each scenario, 40 independent 100 ns simulations were performed with distinct initial velocities and solvent and ion configurations.

### Simulation and network analyses

Equilibrated simulation ensembles were analyzed to determine histidine protonation probabilities, net protein protonation, structural fluctuations, and correlated residue motions. Residue-interaction networks were constructed from persistent structural contacts, electrostatic proximity, and dynamic cross-correlations. Consensus communities were identified using repeated Leiden community detection, and communication within and between communities was quantified from weighted network connections. Node strength and current-flow betweenness centrality were used to identify residues contributing to local connectivity and long-range communication. Statistical comparisons between simulation conditions were performed using permutation testing with false-discovery-rate correction.

## Acknowledgments

We thank to Dr. Eliane Briand for her kind assistance in adapting parameters of the constant-pH molecular dynamics simulations for our system. We also thank Sameer Al-Abdul-Wahid at the NMR centre of the University of Guelph for technical assistance. This research was enabled in part by support provided by the Digital Research Alliance of Canada (alliancecan.ca). This research was supported by funding from NSERC Discovery Grants RGPIN-2022-05139 to E.M.M. and RGPIN-2022-03348 to S.K.; as well as support from the National Institutes of Health through GM R35130587 and S10OD034346 to C.L.B. M.W.-R. acknowledges funding from NSERC PGS-D and the W.S. Rickert Graduate Student Fellowship in Science.

## Author Contributions

I.M.H.M.: Investigation, Validation, Formal analysis, Visualization, Writing - Original draft, Writing - Review & Editing; L.B.P.S.: Investigation, Validation, Formal analysis, Visualization, Writing - Original draft, Writing - Review & Editing; M.W.-R.: Methodology, Formal analysis, Visualization, Writing - Original draft, Writing - Review & Editing; X.L.: Investigation, Methodology, Formal analysis; S.D.: Investigation, Formal analysis; B.G.E.: Resources, Investigation; M.S.B.: Resources; S.K.: Supervision, Formal analysis, Writing - Review & Editing, Funding acquisition; C.L.B.: Conceptualization, Supervision, Resources, Funding acquisition; E.M.M.: Conceptualization, Supervision, Visualization, Writing - Review & Editing, Project administration, Resources, Funding acquisition.

## Competing Interest Statement

The authors declare no competing interest.

## Abbreviations

NMR: nuclear magnetic resonance
CpHMD: Constant-pH Molecular Dynamics
H_(0),88-91_ and H_(0),27-31_: hisactophilin mutant with four histidines substituted for neutral residues in the 88–91 and 27–31 regions, respectively
CSP: chemical shift perturbation
RMSF: root-mean square fluctuation
DCC: dynamic cross-correlation
CFBC: current-flow betweenness centrality

## Notes

### Competing Interest Statement

The authors have declared no competing interest.

