## Supplementary material for "Surface electrostatic networks control hydrophobic core remodeling in a pH-dependent switching protein": SI Appendix

### Supporting Information

**This PDF file includes:**

Supporting text
Figures S1 to S11
Tables S1 to S6
SI References

### Supplementary Methods

#### Mutant design

For studying the roles of surface ionizable residues, we design a set of point and multi-mutans of hisactophilin. These mutants had to meet two general criteria: 1) the residues of interest must be mutated from ionizable residues to non-ionizable residues, and 2) the effect on protein stability should be minimal. In this way, the mutations will alter ionizable residues involved in switching while avoiding introducing other changes to the protein structure or stability. Rational design of mutations involved considering a variety of information sources to guide the selection for residue replacement to decrease the likelihood of selecting an unfavourable mutation, namely:

- A multiple sequence alignment of the fascin proteins to identify residue preferences at each position in hisactophilin, as highly conserved residues tend to be stabilizing[1].
- Residues in structurally equivalent positions of fascins with solved structures to identify alternative residues that are known to exist in the same structural context as the residues of interest in hisactophilin[2].
- Stability predictions were made with the aim of minimizing changes in protein stability coming from the mutations. The stability predictors MAESTRO32[3] and FoldX33[4] were used for this purpose.
- When applicable, beta-turn residue preferences were considered[5].

This rational design led us to the following residue replacements used in this work: H27S; H28A; H30A; H31S; D57G; H88S; H89G; H90G: H91Y; H106S.

#### Protein expression

Mutagenesis was performed by SynBio Technologies (New Jersey, USA) using the custom in-house vector pHW1. Wild type or mutant pHW1 (hisactophilin gene, ampicillin resistance) was transformed into chemically competent BL21 *Escherichia coli* cells carrying the pHV738 plasmid (human N-myristoyltransferase 1 gene, *E. coli* methionylaminopeptidase gene, kanamycin resistance)[6] by heat shock and grown overnight at 37 ºC on LB agar plates containing kanamycin and ampicillin at concentrations of 30 μg/mL and 100 μg/mL, respectively. An isolated colony was used to inoculate a 50 mL LB starter culture (30 μg/mL kanamycin and 100 μg/mL ampicillin) that was grown overnight at 37 ºC in a shaking incubator at 225 rpm. The starter culture was used to inoculate 1 L of M9 minimal media (1:100) containing 30 μg/mL kanamycin and 100 μg/mL ampicillin followed by growth at 37 ºC with shaking at 225 rpm. At $OD_{600\ nm}$ ~0.25, sodium myristate was added to a final concentration of 200 μM. Protein expression was induced at $OD_{600\ nm}$ ~0.7 by addition of IPTG to a final concentration of 1 mM. Expression continued for ~24 hours at 25 ºC with shaking at 225 rpm. Cells were harvested by centrifugation at 5000 × *g* for 15 minutes at 4 ºC. Cells were either lysed immediately or flash frozen in liquid nitrogen and stored at -80 ºC until needed. $^{15}N$-labeled variants for wild type and $H_{(0),88\text{-}91}$ were made by using $^{15}N\text{-}NH_4Cl$ as the sole nitrogen source.

### Cell lysis and protein purification

Cell pellets were resuspended in buffer A (50 mM Tris-HCl, 300 mM NaCl, pH 8) supplemented with 1 mM PMSF and 1 mM $MgCl_2$. DNase I was added to the resuspension and stirred at room temperature until homogenous. Cells were lysed with three passes through an Avestin Emulsiflex-C5 mechanical cell disruptor (Ottawa, Canada) at a pressure >20000 psi. CHAPS was added to the crude lysate at a concentration of 0.5% (w/v) followed by stirring at room temperature for 10 minutes. The lysate was centrifuged at 20000 rpm, 4°C for 30 minutes twice. The soluble fraction was then filtered through a 0.45 µm syringe filter. The clarified lysate was loaded onto a Ni-NTA resin equilibrated with buffer A. After loading, buffer A was run through the column until the $A_{280\ nm}$ returned to baseline. The resin was washed with buffer B (50 mM sodium phosphate, 300 mM NaCl, pH 6.3) for 1 hour or until the $A_{280\ nm}$ returned to baseline. Bound hisactophilin was eluted using buffer C (50 mM sodium acetate, 300 mM NaCl, pH 4). The protein was dialyzed against 4 L of MilliQ water for four hours at room temperature three times. The protein was then concentrated to a volume ~3 mL in an Amicon stirred cell (Millipore Sigma). Myristoylated and non-myristoylated hisactophilin was separated by reverse-phase HPLC (Waters Corporation) on a µBondapak C18 column (Waters Corporation) using a linear acetonitrile gradient from 2% to 62% over 60 minutes with a flow rate of 10 mL/min. All solvents used in HPLC contained TFA at a concentration of 0.1% (v/v). Elution of non-myristoylated and myristoylated hisactophilin was monitored by absorbance at 280 nm using a Waters Corporation 996 photodiode array detector. The separated fractions were dialyzed against 4 L of pH 4 MilliQ water for four hours at room temperature twice, followed by two rounds of exchange against 20 mM ammonium carbonate. Non-myristoylated and myristoylated hisactophilin were concentrated in an Amicon stirred cell (Millipore Sigma) to a concentration of 2 mg/mL as determined by absorbance at 280 nm. Aliquots of 2 mg of protein were prepared and lyophilized overnight. Lyophilized protein was either used immediately or stored at -80 ºC for future use. Mutations and myristoyl group incorporation were verified by electrospray mass spectrometry on a ThermoFisher Q Exactive Oribitrap mass spectrometer. Consistency of the global structure of the mutants was evaluated by inspection of NMR data, confirming minimal deviations with respect to wild type (Fig. S1).

### Chemical denaturation

Lyophilized protein was redissolved to a concentration of 45 µM in 500 mM MES pH 6.2 or potassium phosphate pH 7.7. The protein stock was diluted ten-fold into Eppendorf tubes containing volumes of water and urea corresponding to the desired denaturant concentration (ranging from 0-9 M), giving a final protein and buffer concentration of 4.5 µM and 50 mM in the prepared samples, respectively. Samples were equilibrated in a 25ºC water bath for at least 3 hours following their preparation. Protein unfolding was monitored through intrinsic tyrosine fluorescence with an excitation wavelength of 277 nm and emission scans from 290–350 nm using a Fluorolog 311 fluorimeter (Horiba, Japan). Accurate denaturant concentrations were measured by refractometry. A binomial extrapolation model was fitted to the denaturation curve data, using fixed $m_1$ and $m_2$ values of 2.03 kcal/mol·M and 0.072 kcal/mol·$M^2$, respectively[7,8]:

$$Y = (Y_N + S_N x) - \left((Y_N + S_N x) - (Y_U + S_U x)\right) \frac{e^{\frac{-(m_1 C_{mid} - m_2 C_{mid}^2) + m_1 x - m_2 x^2}{RT}}}{1 + e^{\frac{-(m_1 C_{mid} - m_2 C_{mid}^2) + m_1 x - m_2 x^2}{RT}}} \quad (1)$$

where Y is the observed signal, $Y_N$ is the native signal, $S_N$ is the slope of the native baseline, $Y_U$ is the unfolded signal, $S_U$ is the slope of the unfolded baseline, x is concentration of denaturant, $C_{mid}$ is the midpoint of unfolding, R is the gas constant ($1.987\times10^{-3}$ kcal/mol·K) and T is temperature (298 K).

The myristoyl group provides differing levels of thermodynamic stabilization in the sequestered and accessible states. In this sense, measuring the free energy of unfolding ($\Delta G_U$) for both the myristoylated and non-myristoylated protein at high and low pH allows calculation of the free energy of switching ($\Delta G_{SW}$) using a thermodynamic cycle approach as follows[8,9]:

$$\Delta G_U = m_1 C_{mid} - m_2 C_{mid}^2$$

$$\Delta G_{SW} = \left(\Delta G_{U,myr,7.7} - \Delta G_{U,non-myr,7.7}\right) - \left(\Delta G_{U,myr,6.2} - \Delta G_{U,non-myr,6.2}\right)$$

Here we used pH 7.7 as high pH condition, where most of the protein populates the sequestered state, and pH 6.2 as low pH condition, were there is a significant fraction of accessible state without compromising stability measurements.

**Amide resonance assignments of mutant $H_{(0),88\text{-}91}$**

$^{15}$N-labelled NMR samples were prepared by dissolving lyophilized protein to a final concentration of 3 mM in 50 mM potassium phosphate pH 6.8, with 10% $D_2O$, 1 mM DSS, and 1 mM DTT. 2D-$^{15}$N,$^{1}$H HSQC, 3D-$^{15}$N-edited NOESY-HSQC, and 3D-$^{15}$N-edited TOCSY-HSQC spectra were acquired for myristoylated $H_{(0),88\text{-}91}$ on a 600MHz Bruker Avance III spectrometer equipped with a cryoprobe at the University of Guelph NMR facility. Experiments were executed at 25ºC and used a mixing time of 150 ms for the $^{15}$N-edited NOESY-HSQC and 60 ms for the $^{15}$N-edited TOCSY-HSQC. $H_{(0),88\text{-}91}$ backbone amide assignments were made using the 3D spectra and the wild type assignments as a starting point[7]. Spectra were analyzed using CARA v1.9.1.7.

**NMR monitored pH titration**

Lyophilized protein was redissolved in 50 mM potassium phosphate, pH ~9 to a concentration of 8 mg/mL (~0.6 mM) with 10% $D_2O$ and 1 mM DSS. 1D-$^{1}$H spectra were measured on a either a Bruker Avance 600 MHz spectrometer equipped with a TXI probe at 298 K or a Bruker Avance 700 MHz spectrometer equipped with a cryoprobe. Changes in pH were made using HCl and NaOH 0.2 M stocks. For each pH adjustment, the protein sample was transferred from the NMR tube to an Eppendorf tube using an extended glass pipette. P1 pulse widths were remeasured after every pH adjustment. For 2D-$^{15}N$,$^{1}H$ HSQC pH titration experiments, $^{15}N$-labelled NMR samples of myristoylated wild type and $H_{(0),88\text{-}91}$ were prepared as described above and at a starting pH of ~9. A 1D-$^{1}H$ spectrum was acquired prior to each HSQC, and resonance assignments were transferred and propagated to subsequent pHs by inspection.

Chemical shifts of the myristoyl $CH_3$ and the δ-$CH_3$ of I85 sidechain were obtained using TopSpin v4.3.0, and a pH titration Hill equation was fitted to the data as:

$$\delta = \frac{\delta_{seq} + \delta_{acc}10^{n(pK_{app}-pH)}}{1 + 10^{n(pK_{app}-pH)}} \quad (2)$$

where $\delta_{seq}$ and $\delta_{acc}$ are the chemical shifts of the pure sequestered and accessible states, respectively, $pK_{app}$ is the apparent $pK_a$ value for the switching process, and n is the Hill's coefficient. In addition, chemical shift perturbation (CSP) for the amides group of wild type and $H_{(0),88\text{-}91}$ was calculated as:

$$CSP = \sqrt{\frac{\left({}^{1}H\delta - {}^{1}H\delta_{ref}\right)^2 + \left(\frac{{}^{15}N\delta - {}^{15}N\delta_{ref}}{5}\right)^2}{2}} \quad (3)$$

where $\delta_{ref}$ corresponds to the chemical shift value at pH ~8, used as reference. Pearson correlations between amides' CSP and myristoyl $CH_3$ were calculated and only cases with $p < 0.05$ and $R > 0.97$ are considered.

Amide chemical shift correlations analysis was performed for the assigned residues by using the combined $^{1}H$ and $^{15}N$ chemical shift, calculated as $\delta = 0.2 \times {}^{15}N\delta + {}^{1}H\delta$[10,11]. Dendrograms were constructed via agglomerative clustering by using the pairwise absolute Pearson correlation value between the combined chemical shift of the NHs as a distance parameter ($d = 1 - |R_{ij}|$). Clusters with $|Rij| > 0.97$ are colored as significant in Fig. S3c[11,12].

**Potentiometric measurement for proton binding**

The number of bound proton as a function of pH for myristoylated and non-myristoylated variants of wild type and $H_{(0),88\text{-}91}$ was estimated by potentiometric titration[13]. Lyophilized protein was dissolved to a final concentration of 1 mg/mL (determined by $A_{280\ nm}$) and pH adjusted to a starting value of ~9, ensuring minimal protonation of titratable groups. The solution was then titrated stepwise with small aliquots (1-10 μL) of standardized HCl (0.05 M) while continuously monitoring the pH using a calibrated glass electrode at 25ºC. An identical titration was performed on a blank solution containing all components except the protein. At each pH point, the difference in the cumulative amount of added acid between the protein sample and the blank was attributed to proton binding by the protein. The number of bound protons was estimated from this difference and normalized by the total protein concentration to yield the average number of protons bound per protein molecule (H/protein) as a function of pH.

### Constant pH Molecular Dynamics simulations

The starting structure corresponding to the sequestered conformational state of myristoylated hisactophilin was obtained from previously performed umbrella sampling simulations, which used a model of the structure based on NMR data[7]. We solvated this structure with TIP3P water molecules in a cubic box with a side length of ~68 Å. The system was neutralized and brought to a concentration of 50 mM NaCl with the MMTSB toolset[14]. Next, we computed $pK_a$ values of all titratable residues in the initial structure using PROPKA[15] and assigned protonation states according to the desired pH (6.2 or 7.7). The protein was simulated in the NPT ensemble at 298.15 K and 1 atm for ~1 μs at each condition using pyCHARMM[16]. We distinguish between sequestered and accessible conformational states using our previously defined umbrella sampling coordinate[7]: the distance between the center of the $C_{13}$–$C_{14}$ carbons of the myristoyl group and the centroid of the Cα atoms of Val21, Val61, and Val101 (3V). At pH 7.7, the myristoyl group remained sequestered (distance less than 1.0 nm); however, at pH 6.2 we observed a spontaneous conformational transition from the sequestered state to the accessible state (distance higher than 1.5 nm). From these simulations, we extracted all snapshots where the myristoyl group was accessible in the pH 6.2 simulation and sequestered in the pH 7.7 simulation. Then, we computed the ensemble-averaged $pK_a$ values of all titratable residues for both the sequestered and accessible states using PROPKA[15]. For the second iteration of standard MD simulations, we selected a representative starting structure of the accessible state at pH 6.2 and the sequestered state at pH 7.7 from the first iteration. Additionally, the protonation states were recalculated based on the ensemble-averaged $pK_a$ and the desired pH value. The second iteration of standard MD simulations was run for 1 μs, from which a representative accessible structure at pH 6.2 and sequestered structure at pH 7.7 were obtained and selected for further simulations.

Given the central importance of protonation in the switching mechanism, we employed constant pH molecular dynamics (CpHMD) simulations which allow histidines to dynamically sample their protonation (tautomeric) states in response to the local electrostatic environment. These simulations were performed using a modified GROMACS distribution[17,18] that incorporates fast multipole method (FMM) electrostatics[19,20]. This implementation employs Hamiltonian interpolation through continuous λ-dynamics (where $\lambda_p$ and $\lambda_t$ represent the protonation and tautomeric coordinates, respectively) together with dynamic barrier and well optimization (DBO), which enhances sampling of the protonation states of titratable residues while ensuring that the $\lambda_p$ values primarily occupy the physical end states.

CpHMD simulations were conducted for both sequestered and accessible conformations at two pH conditions, pH 6.2 and pH 7.7, resulting in four independent simulation "scenarios". Each protein structure was placed in an $8 \times 8 \times 8$ $nm^3$ cubic simulation box. All 31 histidine residues were treated as titratable sites, while glutamic and aspartic acids were left deprotonated. CHARMM-modified TIP3P water model[21], sodium, and chloride ions were added to neutralize the system while maintaining a physiological salt concentration of 150 mM. Buffer water molecules required for the CpHMD protocol were added during solvation using the parameters recommended by the method developers (–rminprotld 1.5 nm, –rminld 3.0 nm, and –rtolld 0.3 nm).

Following system setup, energy minimization for up to 50,000 steps was carried out with the steepest descent algorithm to remove unfavorable contacts. This was followed by λ-equilibration for 50 ps, during which all heavy atoms were position-restrained with a force constant of 1000 kJ/molnm$^2$ to allow the λ variables to stabilize. Subsequently, structural equilibration in the NVT ensemble was performed for 500 ps with no restraints. Production simulations were then carried out for 100 ns in the NVT ensemble, since pressure coupling is not currently implemented in this GROMACS distribution.

For each simulation scenario, 40 independent replica simulations of 100 ns each were performed with distinct initial velocities and different ion and water placements to enhance conformational sampling. All simulations employed the CHARMM36m force field[21]. The temperature was maintained at 298 K using the Bussi–Parrinello–Donadio thermostat[22] with a coupling constant of 0.1 ps. A time step of 2 fs was used for integration, with LINCS constraints[23] applied to bonds involving hydrogen atoms. The λ values were recorded every 0.5 ps, while atomic coordinates were saved every 10 ps.

**Constant pH Molecular Dynamics analysis**

The first 50 ns of each trajectory was considered equilibration and discarded based on reverse-cumulative averaging analysis of protein backbone RMSD and $R_g$, the number of hydrogen bonds and contacts, and net protonation, all calculated using MDTraj (version 1.11.1)[24]. The distribution of these metrics in the last 50 ns converged among replicates. As mentioned above, we use our previously defined umbrella sampling coordinate ($M_{C13\text{-}C14}$–3V distance) for defining sequestered and accessible states[7]. Clustering of this distance was performed with Hierarchical Density-Based Spatial Clustering of Applications with Noise (HDBSCAN)[25] in Scikit-learn[26] using a minimum cluster size of 10,000 and a minimum samples parameter of 500. The resulting clusters were used to classify frames as sequestered, accessible, or an intermediate state (with modes of ~0.7 nm, ~2.1 nm and ~1.4 nm, respectively). Analyses were performed on the cluster corresponding to the starting conformation (sequestered or accessible), which was also the dominant state in each scenario, thereby eliminating the contributions from alternate myristoyl conformational states.

Protonation probability (Fig. S4 and Fig. 4b in the main text) was calculated by dividing the number of frames in which a histidine is protonated by the total number of frames, while net protonation is calculated by adding the $\lambda_p$ for all histidines in a single frame and then averaging over all frames. We consider a histidine residue to be protonated (charge of +1) when $\lambda_p$ is higher than 0.5 at that frame and deprotonated otherwise (no charge)[17]. Community net charge (Fig. 5a–d) was calculated by combining the net histidine protonation with the constant charges of other groups (treated as non-ionizable, namely, –1 for aspartate, glutamate, and the C-terminal; +1 for arginine and lysine). For Root Mean Square Fluctuation (RMSF) and Dynamic Cross-Correlation

(DCC) analyses, protein residues were represented by the geometric center of their heavy atoms, and myristoyl was represented by five equally spaced carbon atoms ($C_2$, $C_5$, $C_8$, $C_{11}$, $C_{14}$). RMSF was used to identify regions of increased flexibility or instability while DCC analysis was used to quantify correlated motions between residue geometric centers and selected myristoyl carbon atoms.

To gain a deeper view into the coordinated motions of hisactophilin and identify potential allosteric residues, we constructed two-dimensional interaction networks using NetworkX[27]. Each residue was represented by the geometric center of its heavy atoms, which effectively captures both backbone and side chain dynamics essential for identifying allosteric communication[28], while the myristoyl was represented by the five carbon atoms, as described above. Edges are placed based on the union of two contact definitions: (1) two residues (that are not within ±2 in primary sequence) have heavy atoms within 0.45 nm for >75 % of the sampled frames; and (2) the ionizable atoms of histidine, arginine, lysine, glutamate, aspartate were within 1.0 nm of each other for >75 % of the sampled frames. Edge weights were assigned based on the absolute pairwise DCC value between the connected nodes. Unlike the other charged residues, histidine's ionic state is dynamically sampled and does not consistently satisfy the second contact definition based on electrostatic interactions. Accordingly, edges that only satisfy the second contact definition are scaled by the protonation probability of the connected histidine(s). This analysis provides a quantitative framework for identifying dynamically coupled residue networks and collective motions in proteins[29]. Such correlated motion networks have been shown previously to define allosteric pathways and functional coupling between distant sites[30,31].

We identified communities of nodes with densely (sparsely) connected internal (external) edges by using the Leiden algorithm[32]. Similar approaches have been used to explain allosteric regulation in protein kinases[33] where effector binding redistributes dynamic residue interactions into distinct functional communities that differentiate the active and inactive states of the enzyme. Due to the stochastic nature of this algorithm, single runs can produce varying partitions. We performed consensus community detection by calculating 300 independent Leiden iterations on the network[34]. Based on the pairwise community co-occupancy probability of nodes (between 0 and 1), we formed a consensus matrix. To determine how well our communities were partitioned, we calculated the global mixing parameter ($\mu$), defined as the mean ratio of a given residue's intercommunity weighted edges and its total weighted edges across all residues. We filtered out all entries $\leq 0.5$ as these were considered unlikely co-occupancies resulting from the stochastic algorithm. Next, the Leiden algorithm was applied to the filtered consensus matrix for 300 independent iterations, yielding a new consensus matrix to be filtered. This step of iterating on the filtered matrix was repeated until co-occupancies within communities became 1 and between communities became 0 (this normally occurred within 1 to 2 steps), indicating well-separated and stable communities. To quantify the extent and magnitude of intercommunity communication, we summed the edge weights of all pairs of nodes connecting two given communities. This style of community analysis has been used to investigate domain-level correlations in allosteric networks [31,35].

Two properties were calculated to identify nodes that are important to the network connectivity. Strength is a measure of local collectivity, calculated by summing all edge weights connected to a given node. Current-flow betweenness centrality (CFBC)[36,37] treats the network as an electrical network, with absolute cross-correlation edge weights considered as conductance and quantifies the current flowing through a given node when all paths between node pairs are possible. Unlike betweenness centrality, which is conventionally used, CFBC is not constrained to the geodesic assumption that networks only communicate through their shortest paths. Accordingly, CFBC provides a more robust and accurate description of allosteric networks[38] and has been employed across diverse protein architectures to identify the communication pathways responsible for signal propagation[38-41].

To determine Strength and CFBC stability within simulation scenarios and enable comparisons between simulation scenarios, we split each of our simulation scenarios into five independent groups of eight replicates each, containing an approximately equal number of frames. Strength and CFBC were performed independently for each group, and each resulting value is reported as a mean ± standard deviation in Tables S5. To test for statistically significant differences in each per-node metric between simulation scenarios, the five group-level means were compared using an exact two-sided permutation test of the difference in means. All values, including near zeros, were kept. By enumerating the null distribution of all possible differences in means and comparing it to the observed difference in means, we calculated the p-value for the null hypothesis that the two values are identical. Since we performed many significance tests, our False-Discovery Rate (FDR) is expected to be high. To account for this, we calculated q-values using the Benjamini-Hochberg FDR correction[42] separately for each metric. We consider $q<0.10$ to be significant, meaning that we accept that approximately 10% of our significant differences appear by chance. No significant distribution differences for these values were found between any of the scenarios using a two-sample Kolmogorov-Smirnov test with SciPy (version 1.17.0)[43], testing the null hypothesis that the distributions are identical and considering a p-value less than 0.10 to be significant.

### Supplementary Tables

**Table S1. Analysis of the pH dependence of myristoyl $CH_3$ chemical shift.** A Hill function (eq. 2) was fitted to the myristoyl methyl chemical shift as a function of pH. Chemical shift of pure accessible ($\delta_{acc}$) was set to the same value for all variants and fit as a shared global parameter.

| variant | parameter | | | | $r^2$ |
|---|---|---|---|---|---|
| | $\delta_{seq}$ (ppm) | $\delta_{acc}$ (ppm) | $pK_{app}$ | n | |
| WT | -0.858 ± 0.004 | 0.43 ± 0.06 | 6.00 ± 0.03 | 1.62 ± 0.06 | 0.9948 |
| H106S | -0.851 ± 0.006 | | 5.88 ± 0.03 | 1.48 ± 0.06 | 0.9972 |
| H89G | -0.854 ± 0.006 | | 5.85 ± 0.03 | 1.41 ± 0.07 | 0.9924 |
| H91Y | -0.857 ± 0.006 | | 5.70 ± 0.04 | 1.02 ± 0.04 | 0.9993 |
| D57G | -0.837 ± 0.006 | | 6.16 ± 0.03 | 1.52 ± 0.07 | 0.9987 |
| $H_{(0),88\text{-}91}$ | -0.829 ± 0.004 | | 5.19 ± 0.05 | 0.85 ± 0.03 | 0.9962 |
| D57G/$H_{(0),88\text{-}91}$ | -0.820 ± 0.006 | | 5.45 ± 0.04 | 1.00 ± 0.05 | 0.9978 |
| $H_{(0),27\text{-}31}$ | -0.839 ± 0.004 | | 6.08 ± 0.03 | 1.47 ± 0.05 | 0.9932 |

**Table S2. Equilibrium urea denaturation data for myristoylated and non-myristoylated hisactophilin variants.** A binomial extrapolation model (eq. 1) was fitted to the data with fixed values of $m_1$ and $m_2$ of 2.03 kcal/mol·M and 0.072 kcal/mol·$M^2$, respectively[7]. $C_{MID}$ is the urea concentration at the midpoint of unfolding.

| variant | $C_{MID}$ (M) | | | |
|---|---|---|---|---|
| | pH 6.2 | | pH 7.7 | |
| | myr | non-myr | myr | non-myr |
| WT | 2.96 ± 0.01 | 2.24 ± 0.01 | 5.50 ± 0.01 | 3.92 ± 0.01 |
| H106S | 2.55 ± 0.02 | 1.91 ± 0.02 | 4.96 ± 0.02 | 3.53 ± 0.02 |
| H89G | 2.49 ± 0.01 | 1.90 ± 0.01 | 5.30 ± 0.01 | 3.95 ± 0.01 |
| H91Y | 3.31 ± 0.02 | 2.70 ± 0.02 | 5.63 ± 0.02 | 4.27 ± 0.03 |
| D57G | 2.40 ± 0.01 | 1.86 ± 0.01 | 4.80 ± 0.02 | 3.42 ± 0.01 |
| $H_{(0),88\text{-}91}$ | 3.64 ± 0.03 | 2.75 ± 0.03 | 6.13 ± 0.03 | 4.85 ± 0.03 |
| D57G/$H_{(0),88\text{-}91}$ | 3.37 ± 0.02 | 2.45 ± 0.02 | 5.63 ± 0.04 | 4.22 ± 0.04 |
| $H_{(0),27\text{-}31}$ | 3.23 ± 0.01 | 2.64 ± 0.01 | 5.80 ± 0.02 | 4.30 ± 0.01 |
| **variant** | **$\Delta G_U$ (kcal/mol)** | | | |
| | **pH 6.2** | | **pH 7.7** | |
| | **myr** | **non-myr** | **myr** | **non-myr** |
| WT | 5.38 ± 0.02 | 4.19 ± 0.02 | 8.98 ± 0.02 | 6.86 ± 0.02 |
| H106S | 4.70 ± 0.03 | 3.62 ± 0.03 | 8.30 ± 0.03 | 6.27 ± 0.03 |
| H89G | 4.61 ± 0.02 | 3.59 ± 0.02 | 8.74 ± 0.02 | 6.90 ± 0.02 |
| H91Y | 5.92 ± 0.03 | 4.96 ± 0.03 | 9.14 ± 0.02 | 7.35 ± 0.04 |
| D57G | 4.45 ± 0.02 | 3.53 ± 0.02 | 8.08 ± 0.02 | 6.10 ± 0.02 |
| $H_{(0),88\text{-}91}$ | 6.43 ± 0.05 | 5.05 ± 0.04 | 9.74 ± 0.03 | 8.15 ± 0.04 |
| D57G/$H_{(0),88\text{-}91}$ | 6.03 ± 0.03 | 4.54 ± 0.03 | 9.14 ± 0.05 | 7.28 ± 0.06 |
| $H_{(0),27\text{-}31}$ | 5.81 ± 0.02 | 4.86 ± 0.02 | 9.35 ± 0.02 | 7.39 ± 0.02 |

**Table S3. Analysis of the pH dependence of the I85 δ-$CH_3$ chemical shift.** A Hill function (eq. 2) was fitted to the chemical shift of the I85 δ-methyl as a function of pH.

| variant | parameter | | | | $r^2$ |
|---|---|---|---|---|---|
| | $\delta_{seq}$ (ppm) | $\delta_{acc}$ (ppm) | $pK_{app}$ | n | |
| myr WT | -0.667 ± 0.001 | -0.794 ± 0.002 | 5.86 ± 0.01 | 2.50 ± 0.10 | 0.9939 |
| non-myr WT | -0.702 ± 0.002 | -0.872 ± 0.002 | 6.83 ± 0.02 | 0.98 ± 0.04 | 0.9980 |
| myr $H_{(0),88-91}$ | -0.630 ± 0.001 | -0.700 ± 0.300 | 4.40 ± 0.60 | 0.40 ± 0.05 | 0.9660 |
| non-myr $H_{(0),88-91}$ | -0.737 ± 0.004 | -0.831 ± 0.004 | 6.82 ± 0.05 | 0.79 ± 0.09 | 0.9973 |
| myr D57G/$H_{(0),88-91}$ | -0.621 ± 0.001 | -0.700 ± 0.300 | 4.90 ± 0.50 | 0.49 ± 0.08 | 0.9868 |
| non-myr D57G/$H_{(0),88-91}$ | -0.721 ± 0.003 | -0.818 ± 0.003 | 6.78 ± 0.05 | 1.00 ± 0.10 | 0.9977 |

**Table S4. Residues in the communities found in hisactophilin in the accessible and sequestered states at pH 6.2.** Consensus communities were identified using the Leiden algorithm over the residues (nodes) of the interaction networks of correlation motions obtained in the simulations via the dynamic cross-correlation analysis. Selected residues are coloured to highlight differences in community membership, for mutations characterized here (H27–31, D57, H88–91, and H106) and previously (F6, V36, L76, I85, L93, I116, I118, green), along with additional hydrophobic residues (I55, F74, V83, F113, light green) discussed in the main text as important in core-electrostatic coupling.

| community | sequestered state | accessible state |
|---|---|---|
| **c1** | L53*, S54, I55*, K59, Q60, V61*, Y62, Y92, I93*, S94, V101*, S102, T103*, K104, H106, H107, D108, H109, D110, T111 | mC2‡, mC8‡, G2, H35, V36*, E37, H39, K42, V43*, A44, L45*, K46, L53*, H68, G69, D70, S72, L73, F74* |
| **c2** | F74*, H75, L76*, E77, H78, H79, K82, V83*, S84, I85*, K86, G87, H88, H89, H90, H91, E105, T112, F113* | E19, S54, I55*, G56, D57, H58, K59, Q60, V61*, Y62, H71, G87, H88, H89, H90, H91, T103*, E105 |
| **c3** | mC5‡, H35, V36*, E37, H39, K42, V43*, A44, L45*, K46, Y52, H66, L67, L73 | A5, F6*, K7, S8, H9, H10, G11, H12, F13, L14*, T23*, H24, T32, F113*, E114, I116 |
| **c4** | mC11‡, R4*, G18, E19, H33, F34*, T47, H48, G50, K51, L63, S64, H65 | Y92, I93*, S94, A95*, V101*, S102, K104, H106, H107, D108, H109, D110, T111 |
| **c5** | S8, G11, H12, F13, T23*, H24, H25, A95*, D96, H97, H98 | A16, G18, A20, T47, H48, G50, K51, Y52, L63, S64, H65, H66, L67 |
| **c6** | mC14‡, L14*, S15, A16, E17, A20, V21*, K22, T32, G99, H100 | H75, L76*, E77, H78, H79, K82, V83*, S84, I85*, K86, T112 |
| **c7** | mC8‡, N3, A5, F6*, K7, H9, H10, E114, E115, I116 | S15, E17, V21*, K22, D96, H97, H98, G99, H100 |
| **c8** | D57, H58, H68, D70, H71, S72 | N3, R4*, H33, F34*, E115, I117, I118 |
| **c9** | H27, H28, D29, H30, H31 | H25, H27, H28, D29, H30, H31 |
| **c10** | - | mC5‡, N38 |

*Core residues.

‡Carbon atoms used for representing the myristoyl moiety.

Communities are well-separated, producing global μ values (see Methods) of 0.31 and 0.16 (0.18 and 0.14) for the accessible and sequestered states at pH 6.2 (7.7). Evidently, the accessible state is more difficult to partition, especially at pH 6.2, indicating a more uniformly distributed interaction network with blurred boundaries.

**Table S5. Strength and current flow betweenness centrality obtained from the interaction networks.** Strength is a measure of local collectivity, calculated by summing all edge weights connected to a given node of the network. CFBC treats the network as an electrical network, with absolute pairwise DCC edge weights considered as conductance, and it quantifies the current flowing through a given node when all paths between node pairs are possible. Values are presented as mean ± standard deviation, calculated by separating all replicates into five independent groups with an approximately equal number of frames, with each group using frames from the last 50 ns of simulations corresponding to the main cluster of each conformation.

| Node | Strength | | | | CFBC | | | |
|---|---|---|---|---|---|---|---|---|
| | accessible | | sequestered | | accessible | | sequestered | |
| | pH 6.2 | pH 7.7 | pH 6.2 | pH 7.7 | pH 6.2 | pH 7.7 | pH 6.2 | pH 7.7 |
| **mC2** | 0.4 ± 0.2 | 0.38 ± 0.07 | - | - | 0.01 ± 0.01 | 0.01 ± 0.01 | - | - |
| **mC5** | 0.24 ± 0.08 | 0.3 ± 0.2 | 0.2 ± 0.1 | 0.28 ± 0.07 | 0.01 ± 0.01 | 0.01 ± 0.01 | 0.01 ± 0.01 | 0.00 ± 0.01 |
| **mC8** | 0.15 ± 0.07 | 0.14 ± 0.05 | 0.3 ± 0.2 | 0.5 ± 0.2 | 0.01 ± 0.00 | 0.01 ± 0.00 | 0.02 ± 0.01 | 0.04 ± 0.02 |
| **mC11** | - | 0.07 ± 0.00 | 0.31 ± 0.07 | 0.4 ± 0.1 | - | 0 ± 0 | 0.01 ± 0.02 | 0.02 ± 0.00 |
| **mC14** | - | - | 0.5 ± 0.2 | 0.53 ± 0.09 | - | - | 0.04 ± 0.01 | 0.04 ± 0.01 |
| **GLY2** | 1.1 ± 0.1 | 1.1 ± 0.4 | - | - | 0.03 ± 0.00 | 0.03 ± 0.01 | - | - |
| **ASN3** | 1.5 ± 0.1 | 1.5 ± 0.2 | 1.3 ± 0.4 | 1.2 ± 0.4 | 0.07 ± 0.01 | 0.07 ± 0.01 | 0.02 ± 0.01 | 0.02 ± 0.01 |
| **ARG4** | 2.44 ± 0.09 | 2.0 ± 0.6 | 0.6 ± 0.2 | 0.4 ± 0.2 | 0.13 ± 0.01 | 0.11 ± 0.01 | 0.02 ± 0.01 | 0.02 ± 0.01 |
| **ALA5** | 2.2 ± 0.5 | 1.9 ± 0.8 | 1.1 ± 0.2 | 1.1 ± 0.5 | 0.09 ± 0.02 | 0.09 ± 0.02 | 0.01 ± 0.00 | 0.04 ± 0.04 |
| **PHE6** | 2.1 ± 0.2 | 1.7 ± 0.3 | 2.7 ± 0.5 | 2.3 ± 0.6 | 0.10 ± 0.01 | 0.09 ± 0.01 | 0.10 ± 0.02 | 0.10 ± 0.02 |
| **LYS7** | 2.3 ± 0.3 | 2.2 ± 0.3 | 2.2 ± 0.3 | 2.0 ± 0.6 | 0.10 ± 0.02 | 0.11 ± 0.01 | 0.11 ± 0.02 | 0.10 ± 0.03 |
| **SER8** | 1.2 ± 0.3 | 1.2 ± 0.2 | 1.0 ± 0.1 | 0.9 ± 0.3 | 0.06 ± 0.01 | 0.07 ± 0.01 | 0.04 ± 0.00 | 0.04 ± 0.01 |
| **HIS9** | 0.43 ± 0.05 | 0.3 ± 0.1 | 0.5 ± 0.2 | 0.05 ± 0.03 | 0.01 ± 0.00 | 0.02 ± 0.01 | 0.04 ± 0.00 | 0.02 ± 0.00 |
| **HIS10** | 0.15 ± 0.04 | 0.01 ± 0.00 | 0.19 ± 0.07 | 0.01 ± 0.00 | 0 ± 0 | 0.00 ± 0.01 | 0.01 ± 0.00 | 0 ± 0 |
| **GLY11** | 0.4 ± 0.2 | 0.4 ± 0.1 | 0.38 ± 0.04 | 0.2 ± 0.2 | 0.01 ± 0.01 | 0.00 ± 0.01 | 0 ± 0 | 0 ± 0 |
| **HIS12** | 1.8 ± 0.3 | 1.4 ± 0.3 | 1.5 ± 0.3 | 1.3 ± 0.4 | 0.11 ± 0.02 | 0.13 ± 0.04 | 0.10 ± 0.01 | 0.09 ± 0.03 |
| **PHE13** | 2.7 ± 0.4 | 2.3 ± 0.5 | 1.3 ± 0.3 | 1.0 ± 0.4 | 0.14 ± 0.04 | 0.13 ± 0.03 | 0.11 ± 0.02 | 0.08 ± 0.05 |
| **LEU14** | 2.15 ± 0.08 | 1.9 ± 0.3 | 2.0 ± 0.5 | 2.1 ± 0.2 | 0.1 ± 0.0 | 0.10 ± 0.01 | 0.17 ± 0.02 | 0.18 ± 0.03 |
| **SER15** | 1.3 ± 0.1 | 1.4 ± 0.1 | 1.4 ± 0.2 | 1.8 ± 0.1 | 0.05 ± 0.01 | 0.06 ± 0.01 | 0.07 ± 0.03 | 0.09 ± 0.01 |
| **ALA16** | 0.4 ± 0.4 | 0.5 ± 0.2 | 1.3 ± 0.3 | 1.4 ± 0.3 | 0.01 ± 0.02 | 0.02 ± 0.01 | 0.05 ± 0.01 | 0.05 ± 0.01 |
| **GLU17** | 0.2 ± 0.1 | 0.19 ± 0.04 | 0.36 ± 0.08 | 0.6 ± 0.2 | 0.02 ± 0.01 | 0.02 ± 0.01 | 0.02 ± 0.00 | 0.04 ± 0.04 |
| **GLY18** | 0.19 ± 0.03 | 0.2 ± 0.1 | 0.21 ± 0.09 | 0.26 ± 0.09 | 0 ± 0 | 0 ± 0 | 0 ± 0 | 0 ± 0 |
| **GLU19** | 0.3 ± 0.2 | 0.4 ± 0.1 | 0.6 ± 0.2 | 0.7 ± 0.3 | 0.01 ± 0.01 | 0.02 ± 0.01 | 0.02 ± 0.01 | 0.02 ± 0.01 |
| **ALA20** | 0.2 ± 0.2 | 0.3 ± 0.1 | 0.9 ± 0.2 | 1.3 ± 0.3 | 0.01 ± 0.01 | 0.01 ± 0.01 | 0.04 ± 0.02 | 0.06 ± 0.02 |
| **VAL21** | 1.6 ± 0.3 | 1.8 ± 0.2 | 2.0 ± 0.3 | 1.9 ± 0.3 | 0.09 ± 0.02 | 0.11 ± 0.02 | 0.13 ± 0.01 | 0.12 ± 0.02 |
| **LYS22** | 1.8 ± 0.4 | 1.9 ± 0.3 | 2.0 ± 0.1 | 1.8 ± 0.2 | 0.10 ± 0.02 | 0.11 ± 0.01 | 0.12 ± 0.01 | 0.11 ± 0.02 |
| **THR23** | 1.5 ± 0.3 | 1.9 ± 0.7 | 1.7 ± 0.3 | 1.7 ± 0.2 | 0.07 ± 0.01 | 0.11 ± 0.03 | 0.12 ± 0.02 | 0.13 ± 0.01 |
| **HIS24** | 0.5 ± 0.3 | 0.3 ± 0.1 | 0.8 ± 0.2 | 0.7 ± 0.3 | 0.06 ± 0.02 | 0.05 ± 0.02 | 0.08 ± 0.04 | 0.06 ± 0.02 |
| **HIS25** | 0.03 ± 0.02 | 0.04 ± 0.05 | 0.1 ± 0.1 | 0.1 ± 0.1 | 0.01 ± 0.01 | 0.01 ± 0.02 | 0.00 ± 0.01 | 0.02 ± 0.02 |
| **GLY26** | - | - | 0.13 ± 0.00 | 0.12 ± 0.03 | - | - | 0 ± 0 | 0 ± 0 |
| **HIS27** | 0.24 ± 0.05 | 0.05 ± 0.09 | 0.25 ± 0.08 | 0.01 ± 0.00 | 0.03 ± 0.02 | 0.03 ± 0.03 | 0.02 ± 0.01 | 0.02 ± 0.02 |
| **HIS28** | 0.36 ± 0.02 | 0.02 ± 0.00 | 0.35 ± 0.03 | 0.03 ± 0.00 | 0.02 ± 0.01 | 0 ± 0 | 0.02 ± 0.01 | 0 ± 0 |
| **ASP29** | 0.6 ± 0.1 | 0.05 ± 0.01 | 0.64 ± 0.08 | 0.12 ± 0.09 | 0.04 ± 0.00 | 0.04 ± 0.01 | 0.04 ± 0.01 | 0.05 ± 0.01 |
| **HIS30** | 0.3 ± 0.1 | 0.02 ± 0.01 | 0.3 ± 0.1 | 0.03 ± 0.00 | 0.03 ± 0.01 | 0.03 ± 0.01 | 0.03 ± 0.01 | 0.04 ± 0.02 |
| **HIS31** | 0.12 ± 0.08 | 0.2 ± 0.2 | 0.06 ± 0.06 | 0.02 ± 0.05 | 0.04 ± 0.02 | 0.03 ± 0.03 | 0.03 ± 0.02 | 0.02 ± 0.02 |

| | | | | | | | | |
|---|---|---|---|---|---|---|---|---|
| **THR32** | 0.44 ± 0.07 | 0.6 ± 0.3 | 0.9 ± 0.4 | 1.1 ± 0.2 | 0.02 ± 0.00 | 0.02 ± 0.01 | 0.06 ± 0.01 | 0.06 ± 0.01 |
| **HIS33** | 0.8 ± 0.1 | 0.9 ± 0.3 | 0.8 ± 0.2 | 0.9 ± 0.3 | 0.03 ± 0.00 | 0.03 ± 0.02 | 0.07 ± 0.03 | 0.08 ± 0.04 |
| **PHE34** | 2.1 ± 0.3 | 2.1 ± 0.7 | 1.8 ± 0.2 | 1.9 ± 0.1 | 0.14 ± 0.01 | 0.13 ± 0.03 | 0.12 ± 0.01 | 0.13 ± 0.01 |
| **HIS35** | 2.1 ± 0.1 | 2.4 ± 0.4 | 1.3 ± 0.2 | 1.2 ± 0.3 | 0.09 ± 0.01 | 0.09 ± 0.00 | 0.04 ± 0.00 | 0.04 ± 0.01 |
| **VAL36** | 2.4 ± 0.3 | 2.6 ± 0.4 | 2.1 ± 0.3 | 2.1 ± 0.3 | 0.10 ± 0.02 | 0.11 ± 0.01 | 0.07 ± 0.01 | 0.08 ± 0.01 |
| **GLU37** | 1.3 ± 0.3 | 1.4 ± 0.2 | 2.2 ± 0.4 | 1.6 ± 0.4 | 0.05 ± 0.01 | 0.05 ± 0.01 | 0.09 ± 0.00 | 0.07 ± 0.01 |
| **ASN38** | 0.18 ± 0.06 | 0.6 ± 0.3 | 0.39 ± 0.01 | - | 0.00 ± 0.01 | 0.02 ± 0.02 | 0 ± 0 | - |
| **HIS39** | 0.14 ± 0.08 | 0.3 ± 0.1 | 0.2 ± 0.1 | 0.2 ± 0.2 | 0.01 ± 0.00 | 0.01 ± 0.00 | 0 ± 0 | 0 ± 0 |
| **GLY40** | - | - | - | - | - | - | - | - |
| **GLY41** | - | 0.46 ± 0.03 | - | - | - | 0 ± 0 | - | - |
| **LYS42** | 0.6 ± 0.3 | 1.2 ± 0.2 | 0.6 ± 0.1 | 0.7 ± 0.3 | 0.03 ± 0.02 | 0.07 ± 0.01 | 0.03 ± 0.01 | 0.03 ± 0.01 |
| **VAL43** | 1.9 ± 0.2 | 2.1 ± 0.1 | 2.0 ± 0.5 | 1.5 ± 0.2 | 0.10 ± 0.01 | 0.11 ± 0.01 | 0.09 ± 0.01 | 0.08 ± 0.02 |
| **ALA44** | 2.1 ± 0.4 | 2.2 ± 0.2 | 2.2 ± 0.3 | 1.9 ± 0.4 | 0.09 ± 0.01 | 0.08 ± 0.01 | 0.07 ± 0.01 | 0.08 ± 0.02 |
| **LEU45** | 2.3 ± 0.2 | 2.5 ± 0.2 | 2.5 ± 0.3 | 2.4 ± 0.2 | 0.10 ± 0.01 | 0.11 ± 0.01 | 0.14 ± 0.01 | 0.14 ± 0.01 |
| **LYS46** | 1.9 ± 0.2 | 2.0 ± 0.2 | 2.6 ± 0.7 | 2.2 ± 0.2 | 0.09 ± 0.02 | 0.09 ± 0.01 | 0.10 ± 0.02 | 0.09 ± 0.01 |
| **THR47** | 1.6 ± 0.2 | 1.8 ± 0.3 | 2.5 ± 0.2 | 2.6 ± 0.1 | 0.06 ± 0.01 | 0.06 ± 0.02 | 0.10 ± 0.01 | 0.11 ± 0.01 |
| **HIS48** | 0.42 ± 0.06 | 0.5 ± 0.1 | 0.5 ± 0.2 | 0.58 ± 0.07 | 0.01 ± 0.01 | 0 ± 0 | 0.04 ± 0.01 | 0.03 ± 0.01 |
| **CYS49** | - | - | - | - | - | - | - | - |
| **GLY50** | 0.60 ± 0.07 | 0.7 ± 0.1 | 0.8 ± 0.3 | 0.70 ± 0.05 | 0.00 ± 0.01 | 0.01 ± 0.00 | 0.01 ± 0.01 | 0 ± 0 |
| **LYS51** | 2.8 ± 0.3 | 3.0 ± 0.4 | 2.3 ± 0.2 | 2.4 ± 0.2 | 0.11 ± 0.01 | 0.11 ± 0.01 | 0.10 ± 0.01 | 0.10 ± 0.01 |
| **TYR52** | 2.5 ± 0.1 | 2.0 ± 0.3 | 3.0 ± 0.3 | 3.1 ± 0.5 | 0.11 ± 0.02 | 0.09 ± 0.02 | 0.14 ± 0.01 | 0.14 ± 0.01 |
| **LEU53** | 2.2 ± 0.2 | 2.3 ± 0.2 | 2.0 ± 0.2 | 1.9 ± 0.3 | 0.11 ± 0.01 | 0.13 ± 0.01 | 0.13 ± 0.01 | 0.13 ± 0.02 |
| **SER54** | 1.6 ± 0.3 | 1.8 ± 0.4 | 0.7 ± 0.1 | 0.7 ± 0.3 | 0.09 ± 0.01 | 0.10 ± 0.01 | 0.04 ± 0.01 | 0.04 ± 0.01 |
| **ILE55** | 0.6 ± 0.2 | 0.8 ± 0.1 | 1.0 ± 0.1 | 1.1 ± 0.3 | 0.03 ± 0.01 | 0.05 ± 0.01 | 0.09 ± 0.01 | 0.09 ± 0.03 |
| **GLY56** | 1.2 ± 0.1 | 1.2 ± 0.3 | 0.2 ± 0.1 | - | 0.06 ± 0.01 | 0.06 ± 0.01 | 0 ± 0 | - |
| **ASP57** | 0.8 ± 0.2 | 0.5 ± 0.1 | 0.15 ± 0.08 | 0.1 ± 0.2 | 0.02 ± 0.00 | 0.02 ± 0.00 | 0.01 ± 0.01 | 0.01 ± 0.01 |
| **HIS58** | 0.7 ± 0.1 | 0.19 ± 0.04 | 0.3 ± 0.2 | 0.1 ± 0.1 | 0.02 ± 0.01 | 0.01 ± 0.00 | 0.02 ± 0.00 | 0.01 ± 0.01 |
| **LYS59** | 1.0 ± 0.2 | 0.6 ± 0.3 | 0.7 ± 0.2 | 0.9 ± 0.2 | 0.06 ± 0.01 | 0.04 ± 0.02 | 0.05 ± 0.02 | 0.04 ± 0.02 |
| **GLN60** | 0.9 ± 0.2 | 1.3 ± 0.4 | 0.7 ± 0.1 | 0.9 ± 0.3 | 0.04 ± 0.01 | 0.06 ± 0.01 | 0.05 ± 0.01 | 0.05 ± 0.02 |
| **VAL61** | 1.24 ± 0.05 | 1.3 ± 0.2 | 1.5 ± 0.2 | 1.5 ± 0.2 | 0.07 ± 0.01 | 0.08 ± 0.01 | 0.11 ± 0.03 | 0.11 ± 0.02 |
| **TYR62** | 1.1 ± 0.2 | 1.1 ± 0.4 | 0.6 ± 0.1 | 0.5 ± 0.2 | 0.07 ± 0.01 | 0.07 ± 0.02 | 0.03 ± 0.01 | 0.02 ± 0.01 |
| **LEU63** | 2.2 ± 0.2 | 2.4 ± 0.3 | 2.8 ± 0.2 | 3.1 ± 0.3 | 0.12 ± 0.01 | 0.13 ± 0.02 | 0.16 ± 0.01 | 0.17 ± 0.02 |
| **SER64** | 1.3 ± 0.1 | 0.9 ± 0.3 | 1.0 ± 0.2 | 1.2 ± 0.3 | 0.04 ± 0.00 | 0.03 ± 0.01 | 0.03 ± 0.00 | 0.04 ± 0.01 |
| **HIS65** | 0.5 ± 0.1 | 0.5 ± 0.2 | 0.07 ± 0.02 | 0.3 ± 0.2 | 0.01 ± 0.00 | 0.01 ± 0.01 | 0 ± 0 | 0 ± 0 |
| **HIS66** | 0.36 ± 0.05 | 0.3 ± 0.2 | 0.34 ± 0.09 | 0.32 ± 0.09 | 0.01 ± 0.00 | 0 ± 0 | 0.02 ± 0.01 | 0 ± 0 |
| **LEU67** | 0.8 ± 0.3 | 0.7 ± 0.3 | 0.5 ± 0.2 | 0.7 ± 0.3 | 0.03 ± 0.01 | 0.02 ± 0.02 | 0.02 ± 0.01 | 0.04 ± 0.02 |
| **HIS68** | 0.30 ± 0.08 | 0.3 ± 0.2 | 0.2 ± 0.1 | 0.2 ± 0.1 | 0.01 ± 0.00 | 0.01 ± 0.01 | 0.03 ± 0.02 | 0.01 ± 0.01 |
| **GLY69** | 0.6 ± 0.1 | 0.6 ± 0.2 | - | - | 0.02 ± 0.00 | 0.03 ± 0.01 | - | - |
| **ASP70** | 0.6 ± 0.3 | 0.61 ± 0.08 | 0.3 ± 0.1 | 0.1 ± 0.2 | 0.03 ± 0.01 | 0.03 ± 0.01 | 0.02 ± 0.01 | 0.02 ± 0.01 |
| **HIS71** | 0.8 ± 0.2 | 0.9 ± 0.4 | 0.4 ± 0.1 | 0.4 ± 0.2 | 0.04 ± 0.01 | 0.03 ± 0.01 | 0.04 ± 0.01 | 0.04 ± 0.01 |
| **SER72** | 1.6 ± 0.4 | 1.3 ± 0.4 | 0.3 ± 0.2 | 0.5 ± 0.4 | 0.08 ± 0.01 | 0.07 ± 0.02 | 0.03 ± 0.02 | 0.04 ± 0.03 |
| **LEU73** | 1.7 ± 0.4 | 2.0 ± 0.3 | 1.0 ± 0.2 | 0.9 ± 0.2 | 0.08 ± 0.02 | 0.10 ± 0.01 | 0.04 ± 0.01 | 0.04 ± 0.01 |
| **PHE74** | 2.7 ± 0.2 | 3.0 ± 0.3 | 2.5 ± 0.4 | 2.6 ± 0.3 | 0.16 ± 0.00 | 0.19 ± 0.01 | 0.20 ± 0.03 | 0.19 ± 0.01 |
| **HIS75** | 0.7 ± 0.2 | 0.73 ± 0.09 | 1.1 ± 0.3 | 1.1 ± 0.2 | 0.02 ± 0.00 | 0.02 ± 0.00 | 0.03 ± 0.00 | 0.03 ± 0.00 |
| **LEU76** | 1.29 ± 0.08 | 1.4 ± 0.1 | 1.1 ± 0.2 | 1.1 ± 0.1 | 0.06 ± 0.00 | 0.07 ± 0.01 | 0.05 ± 0.01 | 0.04 ± 0.01 |

| | | | | | | | | |
|---|---|---|---|---|---|---|---|---|
| **GLU77** | 1.24 ± 0.06 | 1.2 ± 0.3 | 1.8 ± 0.5 | 1.5 ± 0.3 | 0.05 ± 0.00 | 0.05 ± 0.01 | 0.10 ± 0.01 | 0.09 ± 0.02 |
| **HIS78** | 0.68 ± 0.08 | 0.4 ± 0.1 | 0.7 ± 0.3 | 0.8 ± 0.2 | 0.02 ± 0.00 | 0.01 ± 0.00 | 0.02 ± 0.01 | 0.01 ± 0.01 |
| **HIS79** | 0.49 ± 0.08 | 0.32 ± 0.05 | 0.68 ± 0.07 | 0.43 ± 0.09 | 0.01 ± 0.00 | 0 ± 0 | 0.02 ± 0.01 | 0 ± 0 |
| **GLY80** | - | - | - | - | - | - | - | - |
| **GLY81** | - | - | 0.06 ± 0.00 | - | - | - | 0 ± 0 | - |
| **LYS82** | 1.5 ± 0.3 | 1.27 ± 0.09 | 1.9 ± 0.7 | 2.5 ± 0.7 | 0.07 ± 0.01 | 0.07 ± 0.00 | 0.08 ± 0.02 | 0.12 ± 0.03 |
| **VAL83** | 2.3 ± 0.2 | 1.8 ± 0.4 | 1.9 ± 0.6 | 2.1 ± 0.4 | 0.11 ± 0.01 | 0.10 ± 0.02 | 0.07 ± 0.00 | 0.08 ± 0.01 |
| **SER84** | 1.41 ± 0.07 | 1.5 ± 0.2 | 1.5 ± 0.5 | 1.8 ± 0.2 | 0.04 ± 0.00 | 0.05 ± 0.01 | 0.06 ± 0.01 | 0.08 ± 0.01 |
| **ILE85** | 2.21 ± 0.07 | 2.2 ± 0.1 | 2.7 ± 0.7 | 2.5 ± 0.4 | 0.10 ± 0.01 | 0.11 ± 0.01 | 0.14 ± 0.01 | 0.12 ± 0.01 |
| **LYS86** | 1.7 ± 0.2 | 1.6 ± 0.2 | 1.9 ± 0.5 | 1.6 ± 0.1 | 0.08 ± 0.01 | 0.08 ± 0.00 | 0.11 ± 0.01 | 0.09 ± 0.01 |
| **GLY87** | 1.2 ± 0.2 | 0.9 ± 0.2 | 1.1 ± 0.2 | 0.9 ± 0.3 | 0.05 ± 0.01 | 0.05 ± 0.02 | 0.06 ± 0.01 | 0.04 ± 0.02 |
| **HIS88** | 0.8 ± 0.1 | 0.7 ± 0.2 | 0.5 ± 0.2 | 0.5 ± 0.2 | 0.04 ± 0.00 | 0.04 ± 0.01 | 0.02 ± 0.02 | 0.02 ± 0.02 |
| **HIS89** | 0.4 ± 0.2 | 0.04 ± 0.03 | 0.4 ± 0.1 | 0.04 ± 0.01 | 0.01 ± 0.01 | 0 ± 0 | 0.01 ± 0.00 | 0 ± 0 |
| **HIS90** | 0.9 ± 0.3 | 0.3 ± 0.1 | 0.8 ± 0.3 | 0.3 ± 0.2 | 0.04 ± 0.01 | 0.03 ± 0.01 | 0.03 ± 0.01 | 0.01 ± 0.01 |
| **HIS91** | 2.1 ± 0.1 | 1.5 ± 0.5 | 1.9 ± 0.5 | 1.7 ± 0.3 | 0.10 ± 0.01 | 0.08 ± 0.02 | 0.10 ± 0.01 | 0.10 ± 0.02 |
| **TYR92** | 2.8 ± 0.3 | 2.9 ± 0.5 | 2.5 ± 0.4 | 2.8 ± 0.4 | 0.11 ± 0.01 | 0.12 ± 0.02 | 0.10 ± 0.02 | 0.11 ± 0.01 |
| **ILE93** | 2.4 ± 0.3 | 2.5 ± 0.3 | 2.7 ± 0.3 | 3.0 ± 0.2 | 0.09 ± 0.01 | 0.10 ± 0.02 | 0.14 ± 0.03 | 0.15 ± 0.01 |
| **SER94** | 2.6 ± 0.2 | 2.5 ± 0.3 | 2.0 ± 0.4 | 2.3 ± 0.2 | 0.07 ± 0.00 | 0.08 ± 0.02 | 0.09 ± 0.03 | 0.10 ± 0.01 |
| **ALA95** | 1.4 ± 0.3 | 1.0 ± 0.3 | 1.1 ± 0.2 | 1.4 ± 0.1 | 0.06 ± 0.01 | 0.05 ± 0.01 | 0.07 ± 0.01 | 0.09 ± 0.03 |
| **ASP96** | 0.9 ± 0.2 | 0.4 ± 0.2 | 0.8 ± 0.3 | 0.5 ± 0.2 | 0.05 ± 0.01 | 0.04 ± 0.02 | 0.05 ± 0.02 | 0.04 ± 0.02 |
| **HIS97** | 0.4 ± 0.1 | 0.4 ± 0.2 | 0.4 ± 0.1 | 0.4 ± 0.2 | 0.02 ± 0.01 | 0.02 ± 0.01 | 0.02 ± 0.01 | 0.02 ± 0.00 |
| **HIS98** | 0.4 ± 0.1 | 0.1 ± 0.1 | 0.40 ± 0.08 | 0.08 ± 0.05 | 0.01 ± 0.01 | 0 ± 0 | 0.02 ± 0.00 | 0.01 ± 0.01 |
| **GLY99** | 0.3 ± 0.2 | 0.4 ± 0.2 | 0.5 ± 0.2 | 0.3 ± 0.2 | 0.00 ± 0.01 | 0.02 ± 0.01 | 0.02 ± 0.01 | 0.01 ± 0.01 |
| **HIS100** | 0.4 ± 0.2 | 0.4 ± 0.2 | 0.57 ± 0.06 | 0.4 ± 0.1 | 0.02 ± 0.01 | 0.01 ± 0.01 | 0.04 ± 0.01 | 0.02 ± 0.01 |
| **VAL101** | 2.4 ± 0.2 | 2.3 ± 0.4 | 2.5 ± 0.2 | 2.6 ± 0.4 | 0.13 ± 0.01 | 0.13 ± 0.02 | 0.18 ± 0.02 | 0.17 ± 0.02 |
| **SER102** | 1.3 ± 0.2 | 1.3 ± 0.2 | 1.6 ± 0.1 | 2.0 ± 0.2 | 0.02 ± 0.01 | 0.02 ± 0.01 | 0.07 ± 0.01 | 0.08 ± 0.02 |
| **THR103** | 1.0 ± 0.1 | 1.1 ± 0.6 | 1.1 ± 0.1 | 1.5 ± 0.2 | 0.06 ± 0.01 | 0.07 ± 0.03 | 0.06 ± 0.01 | 0.07 ± 0.01 |
| **LYS104** | 2.7 ± 0.4 | 2.3 ± 0.5 | 1.3 ± 0.3 | 1.4 ± 0.6 | 0.08 ± 0.01 | 0.09 ± 0.02 | 0.06 ± 0.01 | 0.06 ± 0.03 |
| **GLU105** | 1.1 ± 0.3 | 0.4 ± 0.1 | 0.9 ± 0.4 | 0.7 ± 0.4 | 0.05 ± 0.01 | 0.02 ± 0.01 | 0.04 ± 0.01 | 0.04 ± 0.01 |
| **HIS106** | 1.2 ± 0.2 | 0.6 ± 0.2 | 1.1 ± 0.2 | 0.6 ± 0.1 | 0.04 ± 0.01 | 0.01 ± 0.00 | 0.05 ± 0.01 | 0.01 ± 0.00 |
| **HIS107** | 1.3 ± 0.2 | 0.9 ± 0.1 | 1.0 ± 0.3 | 0.8 ± 0.2 | 0.03 ± 0.00 | 0.01 ± 0.00 | 0.03 ± 0.01 | 0.01 ± 0.00 |
| **ASP108** | 2.6 ± 0.4 | 2.0 ± 0.2 | 1.6 ± 0.4 | 1.2 ± 0.5 | 0.06 ± 0.01 | 0.06 ± 0.01 | 0.07 ± 0.01 | 0.05 ± 0.02 |
| **HIS109** | 1.4 ± 0.3 | 0.5 ± 0.1 | 0.7 ± 0.3 | 0.3 ± 0.2 | 0.04 ± 0.01 | 0.02 ± 0.01 | 0.03 ± 0.01 | 0.02 ± 0.00 |
| **ASP110** | 3.1 ± 0.6 | 2.5 ± 0.5 | 1.4 ± 0.4 | 1.4 ± 0.5 | 0.10 ± 0.02 | 0.10 ± 0.02 | 0.07 ± 0.02 | 0.08 ± 0.02 |
| **THR111** | 3.1 ± 0.4 | 3.2 ± 0.4 | 1.5 ± 1.1 | 2.2 ± 0.6 | 0.08 ± 0.00 | 0.1 ± 0.0 | 0.06 ± 0.03 | 0.09 ± 0.02 |
| **THR112** | 1.1 ± 0.3 | 1.1 ± 0.2 | 1.0 ± 0.4 | 1.3 ± 0.2 | 0.04 ± 0.01 | 0.05 ± 0.01 | 0.02 ± 0.01 | 0.04 ± 0.02 |
| **PHE113** | 2.9 ± 0.3 | 2.8 ± 0.4 | 2.6 ± 0.8 | 2.6 ± 0.6 | 0.15 ± 0.02 | 0.17 ± 0.01 | 0.15 ± 0.03 | 0.17 ± 0.01 |
| **GLU114** | 1.53 ± 0.05 | 1.4 ± 0.2 | 2.1 ± 0.2 | 1.9 ± 0.3 | 0.06 ± 0.00 | 0.06 ± 0.01 | 0.08 ± 0.01 | 0.08 ± 0.02 |
| **GLU115** | 1.0 ± 0.1 | 0.8 ± 0.4 | 2.2 ± 0.4 | 2.0 ± 0.7 | 0.05 ± 0.01 | 0.04 ± 0.01 | 0.05 ± 0.01 | 0.05 ± 0.01 |
| **ILE116** | 1.8 ± 0.2 | 1.7 ± 0.4 | 2.2 ± 0.3 | 1.8 ± 0.8 | 0.07 ± 0.01 | 0.08 ± 0.01 | 0.06 ± 0.01 | 0.06 ± 0.02 |
| **ILE117** | 0.46 ± 0.08 | 0.3 ± 0.2 | 0.13 ± 0.00 | - | 0 ± 0 | 0 ± 0 | 0 ± 0 | - |
| **ILE118** | 1.2 ± 0.3 | 0.9 ± 0.6 | - | - | 0.06 ± 0.03 | 0.04 ± 0.02 | - | - |

**Table S6. Differences in Strength and current flow betweenness centrality at equivalent pH obtained from the interaction networks.** Statistically significant differences (q<0.10) are bolded based on an exact two-sided permutation test of the difference in means across the five separated groups of replicates, followed by FDR correction. Raw Strength and CFBC values are listed in Table S5.

| Node | ΔStrength (Seq – Acc) | | ΔCFBC (Seq – Acc) | | Node | ΔStrength (Seq – Acc) | | ΔCFBC (Seq – Acc) | | Node | ΔStrength (Seq – Acc) | | ΔCFBC (Seq – Acc) | |
|---|---|---|---|---|---|---|---|---|---|---|---|---|---|---|
| | pH 6.2 | pH 7.7 | pH 6.2 | pH 7.7 | | pH 6.2 | pH 7.7 | pH 6.2 | pH 7.7 | | pH 6.2 | pH 7.7 | pH 6.2 | pH 7.7 |
| mC2 | - | - | - | - | ASN38 | 0.21 | - | 0.00 | - | HIS79 | **0.19** | 0.10 | **0.01** | **0.00** |
| mC5 | -0.01 | 0.02 | 0.00 | -0.01 | HIS39 | 0.01 | -0.15 | **-0.01** | -0.01 | GLY80 | - | - | - | - |
| mC8 | 0.14 | 0.36 | 0.02 | 0.04 | GLY40 | - | - | - | - | GLY81 | - | - | - | - |
| mC11 | - | 0.36 | - | 0.02 | GLY41 | - | - | - | - | LYS82 | 0.37 | **1.20** | 0.02 | **0.05** |
| mC14 | - | - | - | - | LYS42 | -0.04 | -0.50 | -0.01 | **-0.04** | VAL83 | -0.38 | 0.25 | **-0.04** | -0.01 |
| GLY2 | - | - | - | - | VAL43 | 0.09 | **-0.64** | -0.01 | -0.03 | SER84 | 0.06 | 0.25 | **0.01** | **0.02** |
| ASN3 | -0.27 | -0.25 | **-0.05** | **-0.05** | ALA44 | 0.14 | -0.27 | -0.01 | -0.01 | ILE85 | 0.45 | 0.28 | **0.04** | 0.01 |
| ARG4 | **-1.89** | **-1.59** | **-0.11** | **-0.10** | LEU45 | 0.18 | -0.09 | **0.04** | **0.04** | LYS86 | 0.15 | 0.03 | **0.03** | 0.01 |
| ALA5 | -1.07 | -0.76 | -0.08 | -0.05 | LYS46 | **0.65** | 0.19 | 0.01 | 0.00 | GLY87 | -0.09 | 0.02 | 0.01 | -0.01 |
| PHE6 | 0.55 | 0.55 | 0.01 | 0.01 | THR47 | **0.87** | **0.78** | **0.04** | **0.05** | HIS88 | **-0.31** | -0.19 | -0.02 | -0.02 |
| LYS7 | -0.09 | -0.16 | 0.01 | -0.01 | HIS48 | 0.09 | 0.12 | **0.03** | **0.03** | HIS89 | 0.02 | 0.00 | 0.00 | 0.00 |
| SER8 | -0.29 | -0.29 | **-0.01** | **-0.02** | CYS49 | - | - | - | - | HIS90 | -0.13 | 0.04 | -0.01 | -0.02 |
| HIS9 | 0.05 | **-0.26** | **0.02** | 0.00 | GLY50 | 0.20 | 0.05 | 0.00 | -0.01 | HIS91 | -0.21 | 0.20 | 0.00 | 0.01 |
| HIS10 | 0.04 | 0.00 | 0.00 | 0.00 | LYS51 | -0.49 | **-0.59** | -0.01 | -0.01 | TYR92 | -0.30 | -0.11 | 0.00 | -0.01 |
| GLY11 | -0.04 | -0.14 | -0.01 | 0.00 | TYR52 | **0.49** | **1.05** | **0.03** | **0.05** | ILE93 | 0.30 | 0.45 | **0.05** | **0.05** |
| HIS12 | -0.30 | -0.16 | -0.01 | -0.04 | LEU53 | -0.16 | -0.40 | **0.02** | 0.00 | SER94 | **-0.56** | -0.21 | 0.02 | 0.02 |
| PHE13 | **-1.47** | **-1.33** | -0.04 | -0.05 | SER54 | **-0.88** | **-1.10** | **-0.04** | **-0.05** | ALA95 | -0.26 | 0.33 | 0.01 | **0.05** |
| LEU14 | -0.15 | 0.23 | **0.07** | **0.09** | ILE55 | **0.42** | 0.26 | **0.06** | **0.05** | ASP96 | -0.01 | 0.05 | 0.00 | 0.01 |
| SER15 | 0.16 | **0.46** | 0.02 | 0.02 | GLY56 | -0.92 | - | -0.06 | - | HIS97 | 0.03 | -0.01 | 0.00 | 0.00 |
| ALA16 | **0.86** | **0.92** | **0.04** | **0.03** | ASP57 | **-0.68** | **-0.41** | -0.01 | -0.01 | HIS98 | 0.05 | -0.06 | 0.00 | 0.00 |
| GLU17 | 0.16 | **0.36** | 0.00 | 0.02 | HIS58 | **-0.45** | -0.11 | 0.01 | 0.00 | GLY99 | 0.21 | -0.17 | 0.02 | -0.01 |
| GLY18 | 0.02 | 0.07 | 0.00 | 0.00 | LYS59 | -0.30 | 0.25 | -0.01 | 0.00 | HIS100 | 0.16 | -0.01 | **0.02** | 0.01 |
| GLU19 | 0.29 | 0.30 | 0.01 | 0.01 | GLN60 | -0.24 | -0.34 | 0.01 | 0.00 | VAL101 | 0.09 | 0.36 | **0.04** | **0.04** |
| ALA20 | **0.66** | **0.97** | **0.04** | **0.05** | VAL61 | **0.30** | 0.17 | 0.04 | 0.03 | SER102 | **0.32** | **0.73** | **0.05** | **0.06** |
| VAL21 | 0.33 | 0.12 | **0.04** | 0.00 | TYR62 | **-0.52** | -0.59 | **-0.04** | **-0.05** | THR103 | 0.08 | 0.38 | 0.00 | 0.00 |
| LYS22 | 0.20 | -0.19 | 0.02 | 0.00 | LEU63 | **0.66** | **0.71** | **0.04** | 0.04 | LYS104 | **-1.42** | -0.97 | **-0.02** | -0.02 |
| THR23 | 0.19 | -0.22 | **0.05** | 0.02 | SER64 | **-0.35** | 0.29 | **-0.01** | 0.01 | GLU105 | -0.15 | 0.30 | -0.01 | 0.01 |
| HIS24 | 0.26 | 0.37 | 0.02 | 0.01 | HIS65 | **-0.37** | -0.14 | -0.01 | -0.01 | HIS106 | -0.10 | -0.04 | 0.01 | 0.00 |
| HIS25 | 0.06 | 0.03 | -0.01 | 0.00 | HIS66 | -0.02 | -0.03 | 0.01 | 0.00 | HIS107 | -0.32 | -0.03 | 0.00 | 0.00 |
| GLY26 | - | - | - | - | LEU67 | -0.35 | 0.02 | -0.01 | 0.01 | ASP108 | **-1.08** | **-0.83** | 0.01 | -0.01 |
| HIS27 | 0.00 | -0.04 | -0.01 | -0.01 | HIS68 | -0.06 | -0.15 | 0.02 | 0.00 | HIS109 | **-0.70** | -0.13 | -0.02 | 0.00 |
| HIS28 | -0.01 | 0.01 | 0.00 | 0.00 | GLY69 | - | - | - | - | ASP110 | **-1.67** | **-1.06** | **-0.03** | -0.02 |
| ASP29 | 0.01 | **0.06** | 0.00 | 0.01 | ASP70 | -0.33 | **-0.48** | -0.01 | -0.01 | THR111 | -1.60 | -0.95 | -0.02 | -0.01 |
| HIS30 | 0.02 | 0.01 | 0.00 | 0.01 | HIS71 | **-0.34** | -0.51 | 0.00 | 0.01 | THR112 | -0.11 | 0.15 | -0.02 | -0.01 |
| HIS31 | -0.06 | -0.15 | -0.01 | -0.01 | SER72 | **-1.29** | -0.79 | **-0.05** | -0.03 | PHE113 | -0.29 | -0.23 | 0.01 | 0.00 |
| THR32 | 0.44 | **0.56** | 0.04 | **0.04** | LEU73 | **-0.68** | **-1.11** | **-0.05** | **-0.06** | GLU114 | **0.52** | **0.48** | **0.02** | 0.01 |
| HIS33 | 0.00 | 0.04 | **0.03** | 0.05 | PHE74 | -0.16 | -0.44 | **0.04** | 0.00 | GLU115 | **1.17** | **1.21** | 0.00 | 0.01 |
| PHE34 | -0.28 | -0.13 | -0.02 | 0.00 | HIS75 | **0.43** | **0.35** | **0.02** | **0.01** | ILE116 | **0.44** | 0.13 | -0.01 | -0.02 |
| HIS35 | **-0.81** | **-1.21** | **-0.05** | **-0.05** | LEU76 | -0.17 | **-0.24** | **-0.02** | **-0.03** | ILE117 | -0.32 | - | 0.00 | - |
| VAL36 | -0.28 | -0.51 | **-0.04** | -0.02 | GLU77 | **0.56** | 0.31 | **0.04** | **0.04** | ILE118 | - | - | - | - |
| GLU37 | **0.89** | 0.20 | **0.04** | **0.02** | HIS78 | 0.04 | **0.44** | 0.00 | 0.01 | | | | | |

In principle, Strength and CFBC differences can be calculated for the same, or different, states at different pH ranges. We find that this type of comparison generally produces uninformative trends, primarily highlighting histidines that are changing their protonation state (which we already characterized in Fig. S5 and 4b in the main text).

The magnitude of CFBC values for the surface electrostatic residues is consistently lower than for hydrophobic and core residues (Table S5), indicating that the communication is more efficiently through the network of hydrophobic interactions. Nevertheless, at pH 6.2, the surface electrostatic residues do present significant differences in CFBC (Table S6). Of these significant differences between myristoyl states, about twice as many show increased values in the sequestered state. Together with the larger fraction of predominantly repulsive intercommunity communication (Fig. 5c and d in the main text) and lower surface Strength values (Table S6 and Fig. 4d in the main text), this indicates a more constrained surface communication network in the sequestered state. In contrast, the accessible state has a more locally coupled surface network due to the formation of alternative communication pathways.

### Supplementary Figures

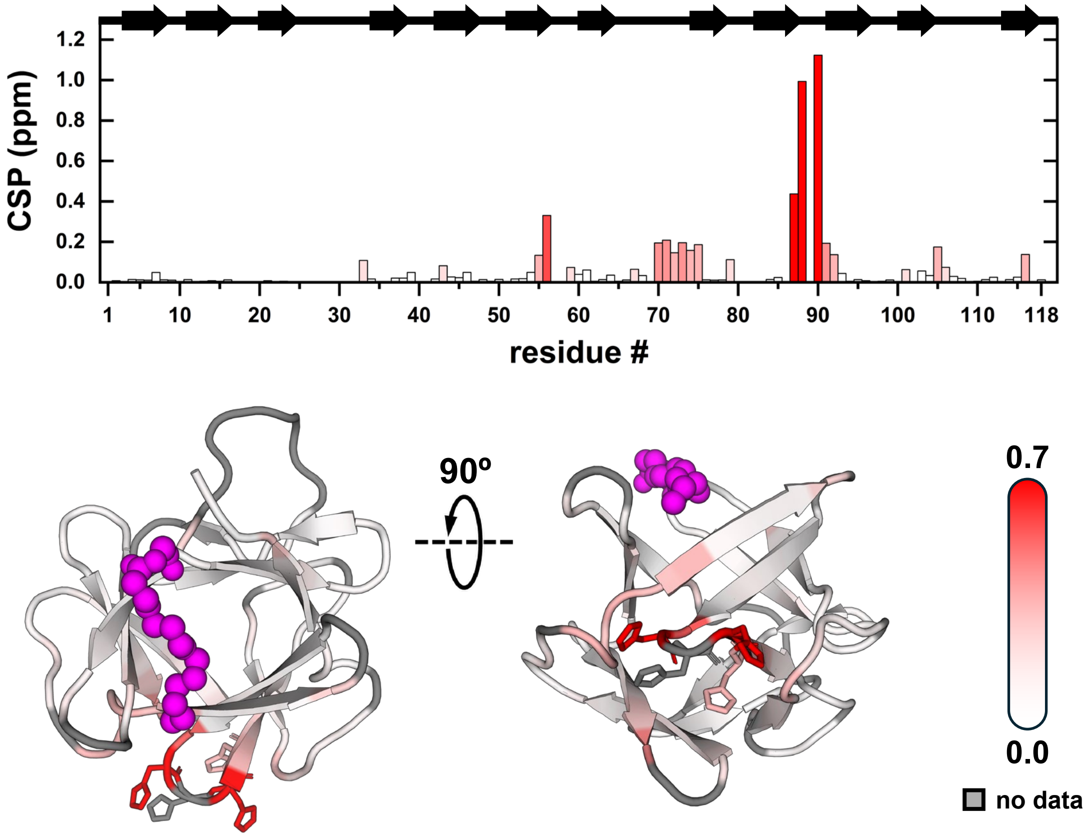

**Figure S1. Chemical shift perturbation of H$_{(0),88\text{-}91}$ mutant with respect to wild type.** CSP is calculated as in eq. 3 using both $^{1}$H and $^{15}$N chemical shift of the mutant and wild type hisactophilin at pH ~8. Residues mutated (H88–91) display the highest CSP values and are shown as sticks in the ribbon representation, while the rest of the protein display small values, signalling that structure is not affected by the mutation.

**Figure S2. pH-dependence of amide chemical shifts for myristoylated wild type and H$_{(0),88\text{-}91}$ mutant.** Plots show the $^{1}$H and $^{15}$N chemical shifts as well as the chemical shift perturbation (CSP) with respect to high pH (~8) condition (eq. 3) for both wild type (black symbols) and H$_{(0),88\text{-}91}$ mutant (purple symbols).

(Plots are shown on the following pages)

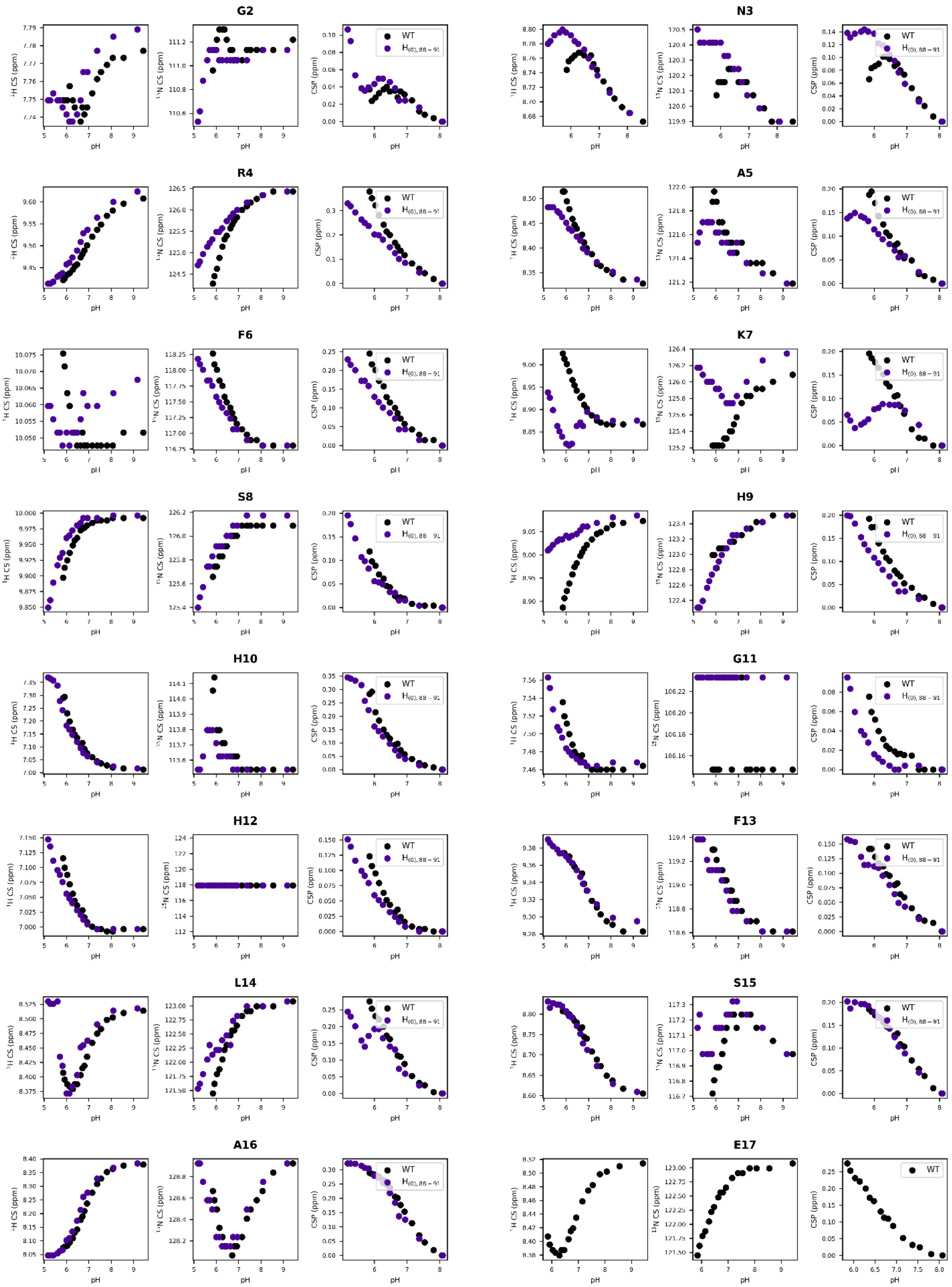

**Figure S2 (cont.).**

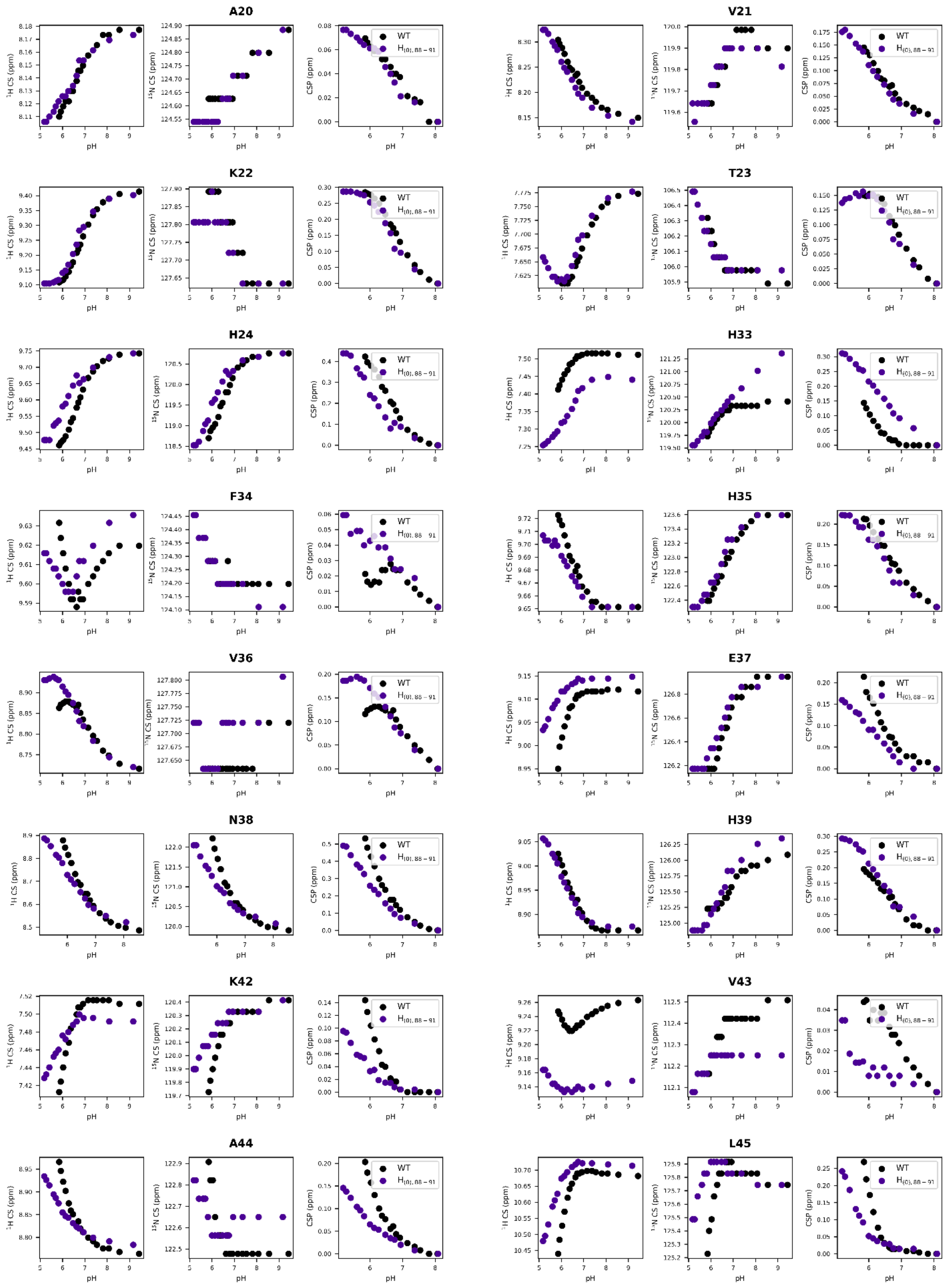

**Figure S2 (cont.).**

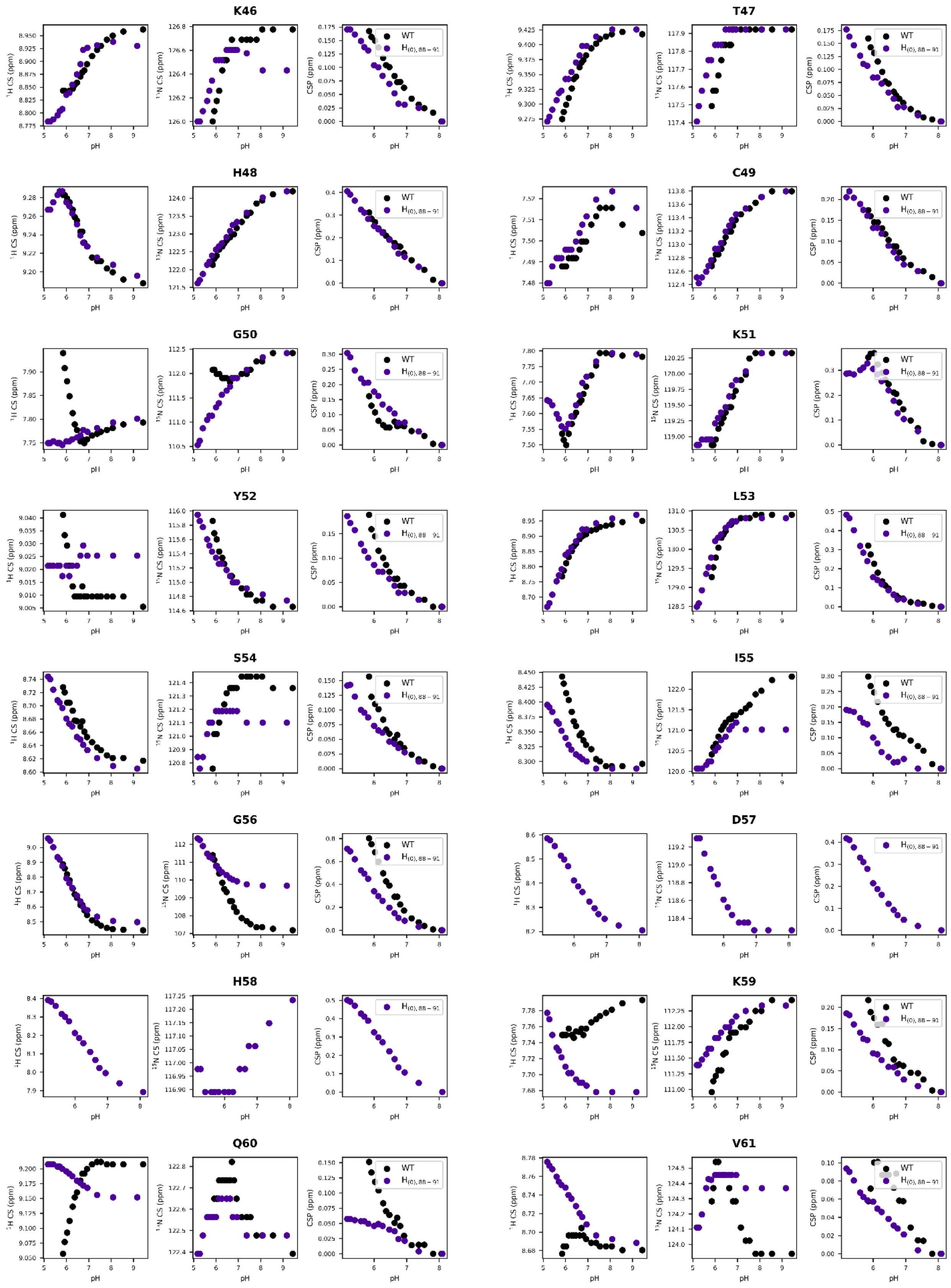

**Figure S2 (cont.).**

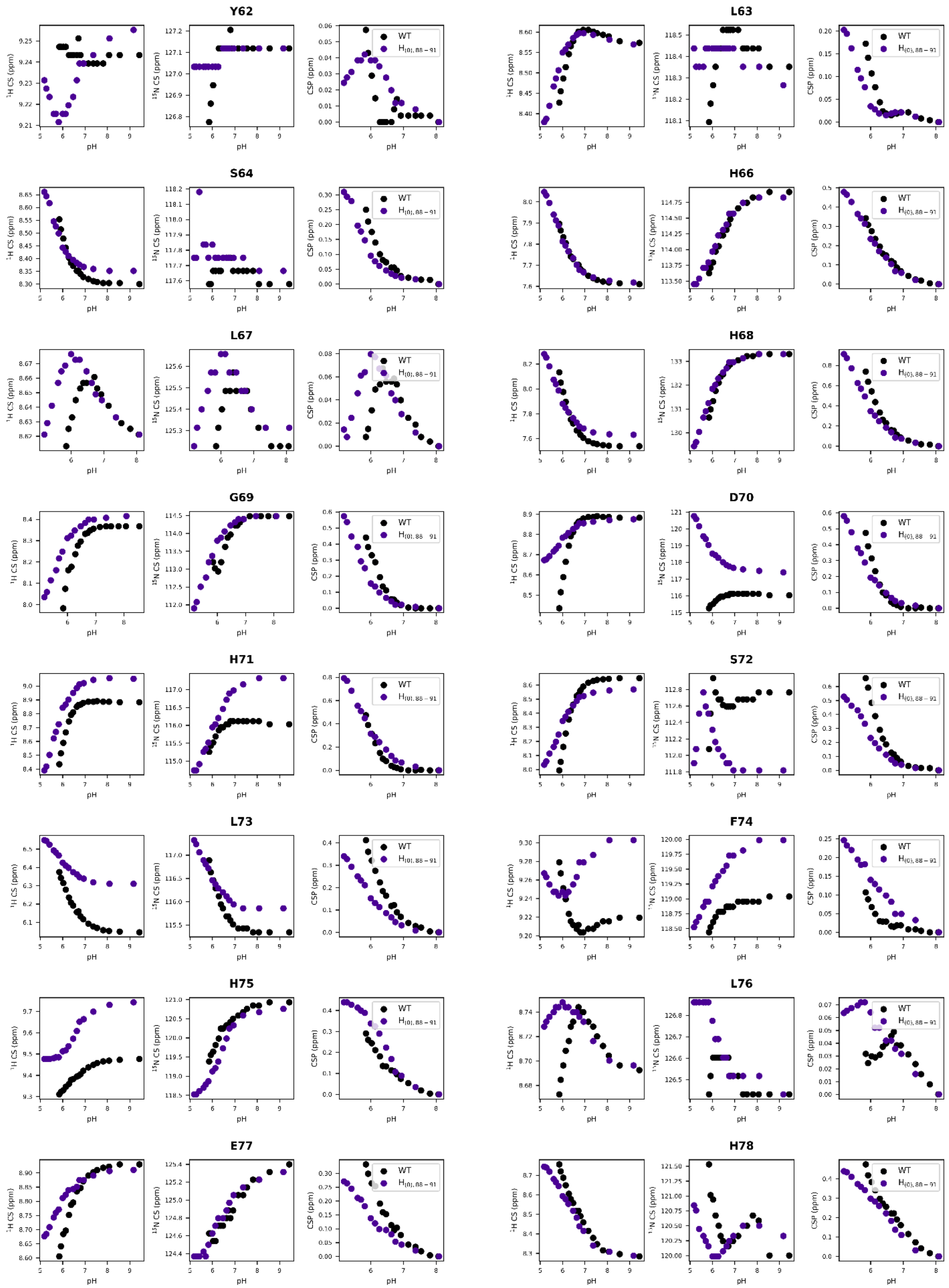

**Figure S2 (cont.).**

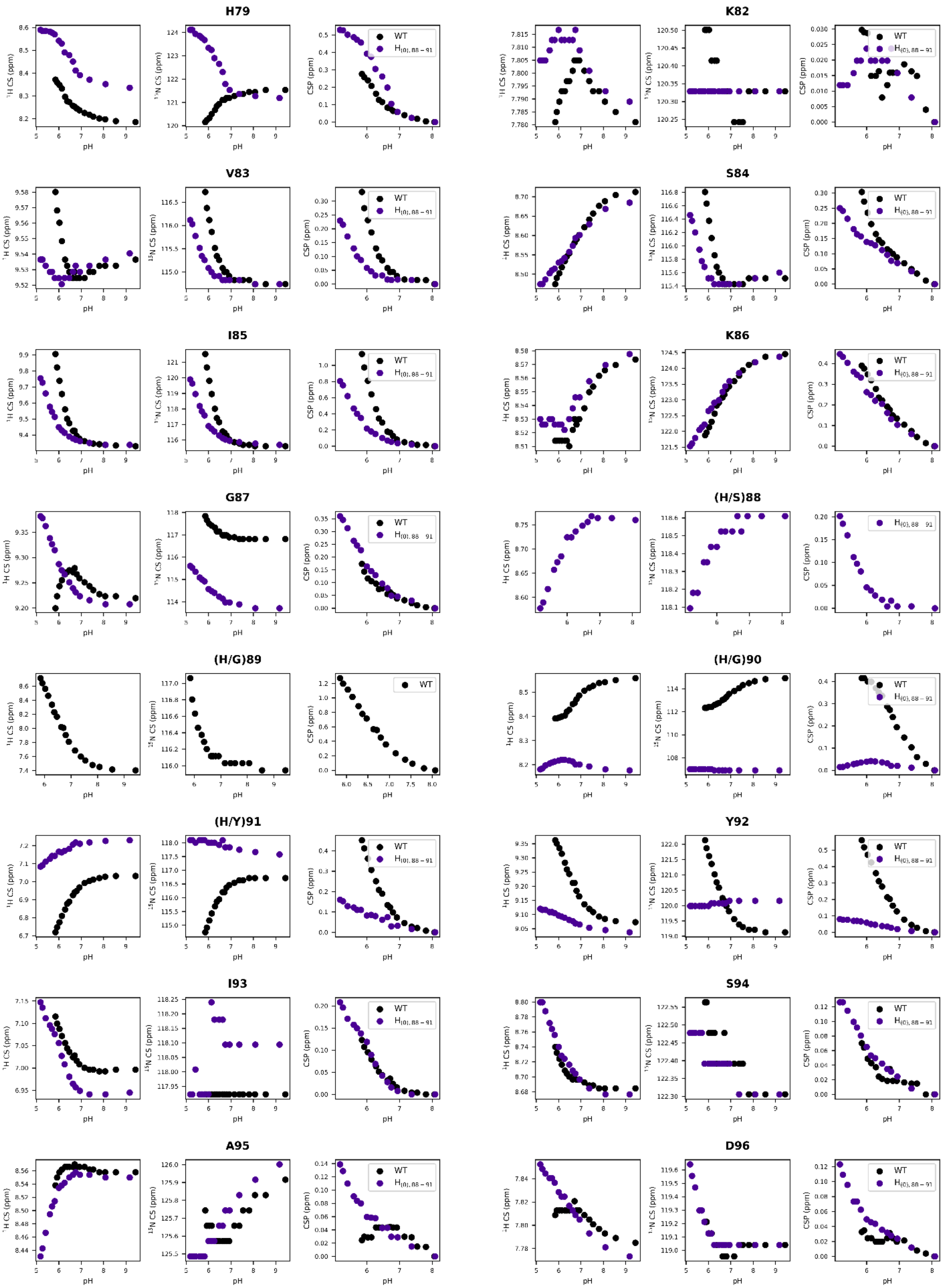

**Figure S2 (cont.).**

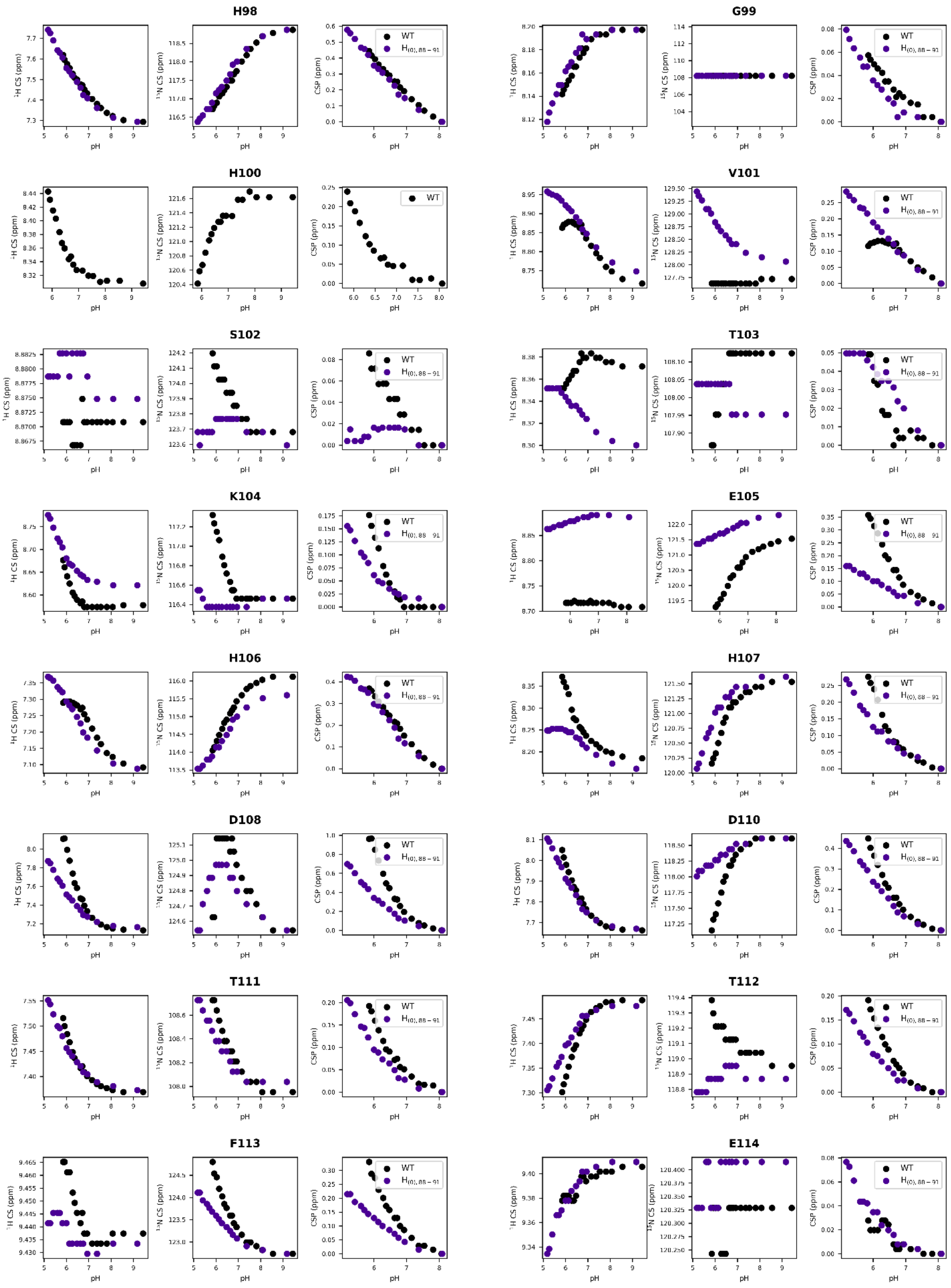

**Figure S2 (cont.).**

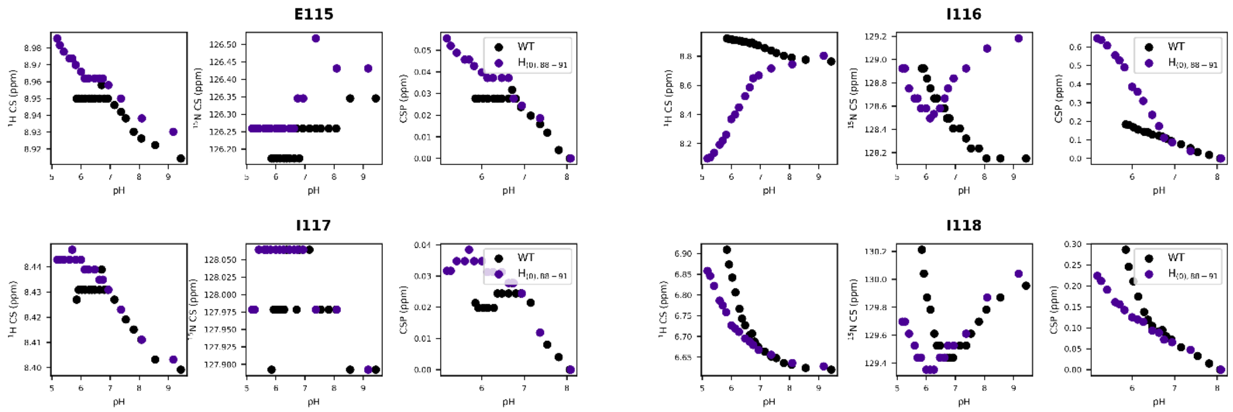

**Figure S2 (cont.).**

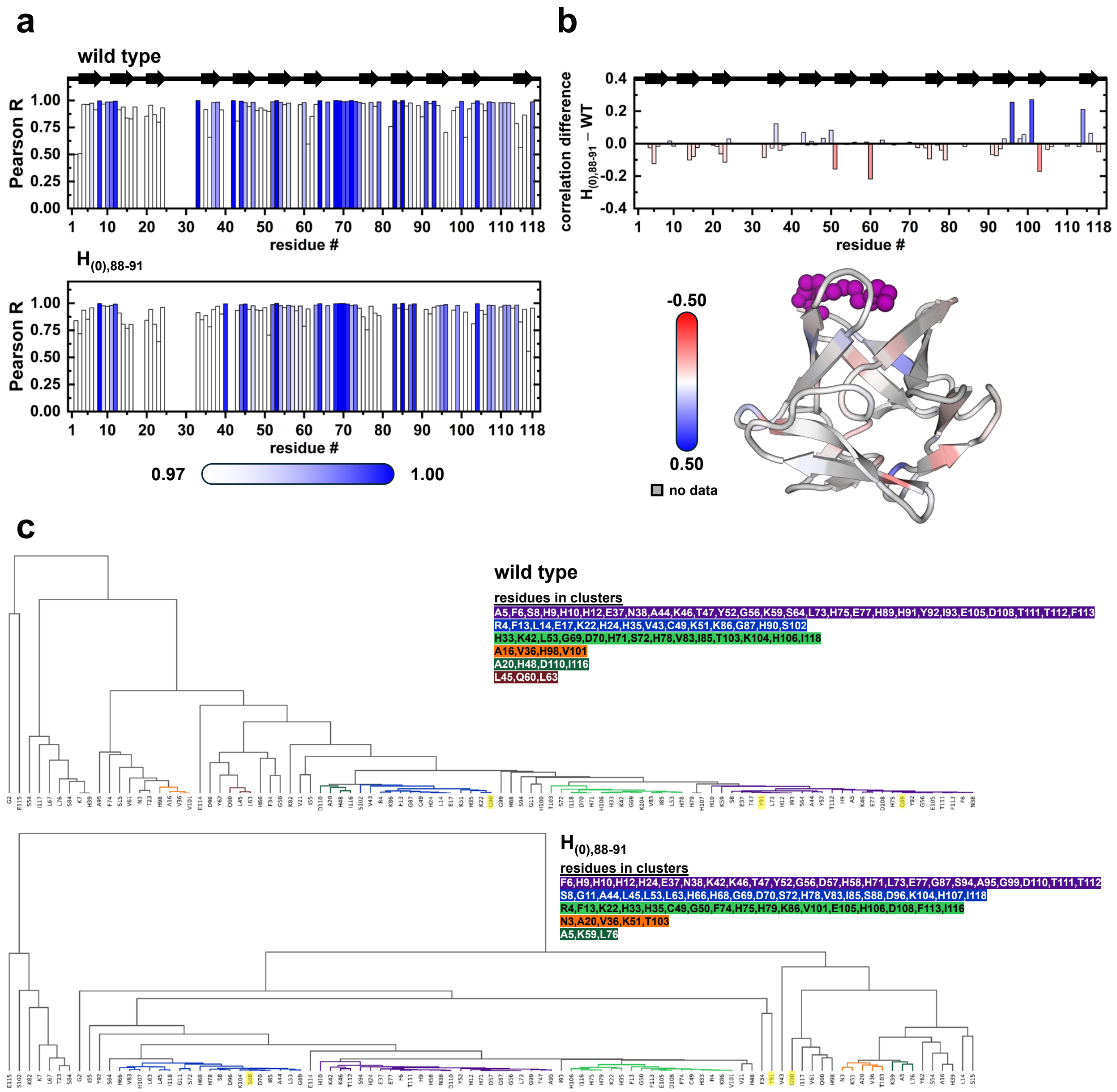

**Figure S3. Analysis of chemical shift correlations for wild type and $H_{(0),88-91}$ mutant HSQC data.** (a) Pearson correlation coefficient between myristoyl $CH_3$ chemical shift and residues' NH chemical shift perturbation (CSP) across the studied range of pH (~5 to ~9, see Figure S1) for wild type (top) and $H_{(0),88-91}$ mutant (bottom). (b) Difference between the Pearson R values of figure (a) as mutant – wild type, represented both in primary sequence (top) and onto the 3D structure (bottom). The difference is not calculated (no data) if a residue is not assigned in either wild type or $H_{(0),88-91}$ mutant. (c) Dendrograms constructed from the pairwise correlation analysis of NH combined chemical shifts[11,12]. Clusters with $|R_{ij}| > 0.97$ and their residues are shown in colors, depicting rearrangement of the networks due to the mutations.

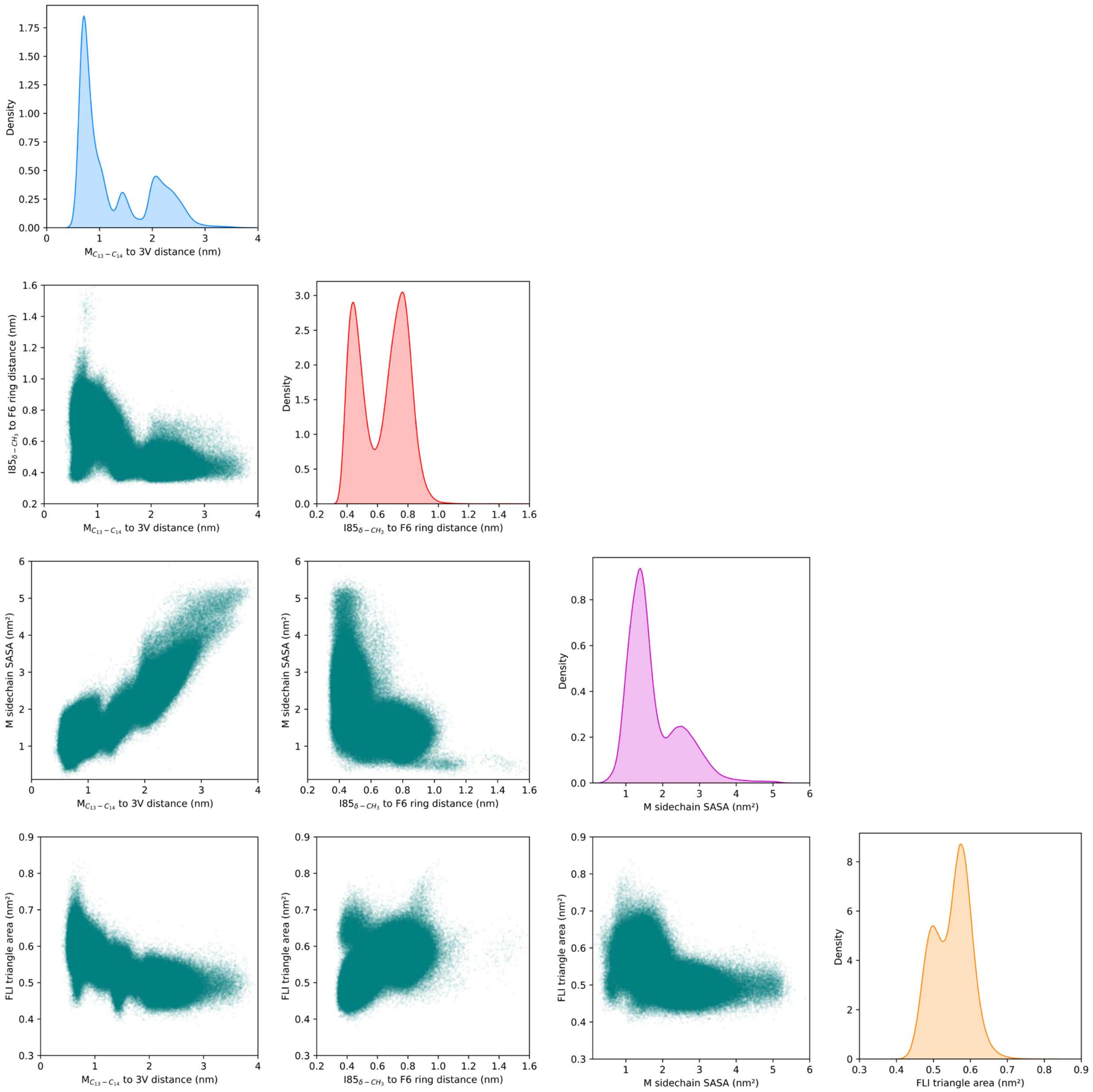

**Figure S4. Distribution of four geometrical parameters obtained in the constant pH molecular dynamics simulations.** Diagonal plots illustrate the density histograms of the four metrics used: distance from the center of the $C_{13}$–$C_{14}$ bond of the myristoyl group to the centroid of the Cα of Val21, Val61, and Val101 (blue); distance from the δ-methyl of I85 to the center of the aromatic ring of F6 (red); solvent accessible surface area (SASA) of the myristoyl acyl chain (magenta); and the area of the FLI triangle, formed by the Cα of residues F6–L45–I85 (orange). Off-diagonal graphs illustrate pairwise scatter plots among the four metrics. Data is obtained from the last 50 ns of the 40 replicates simulations for the four scenarios studied in this work (i.e., starting structure as accessible or sequestered, and at pH 6.2 or 7.7).

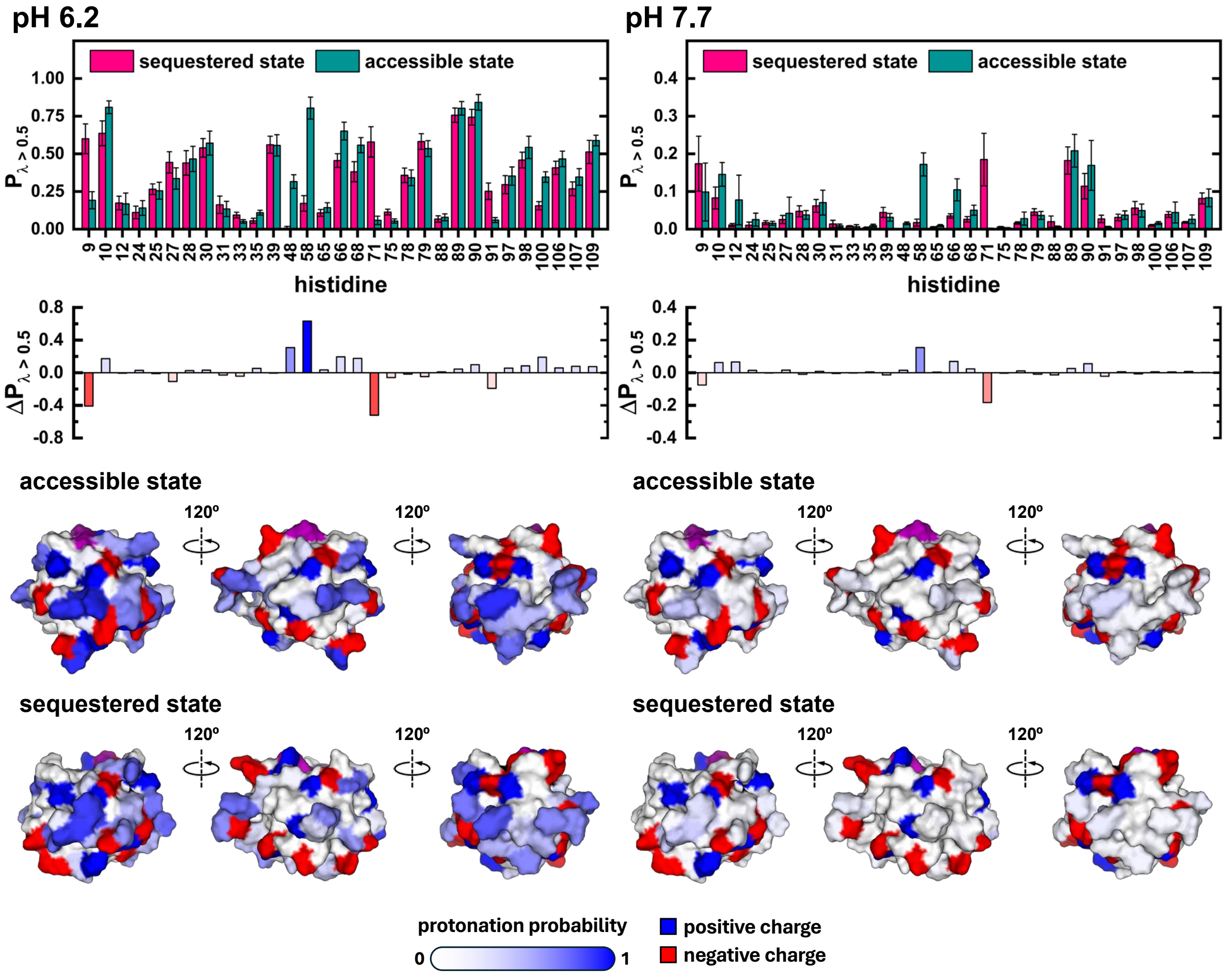

**Figure S5. Histidines protonation probability obtained from constant pH molecular dynamics simulations.** Top bar graphs illustrate the protonation probability for histidine residues in sequestered (pink) and accessible (dark cyan) states at pH 6.2 (left panel) and 7.7 (right panel). The difference in protonation probability is depicted in the bars at the bottom of each panel, calculated as accessible – sequestered. Probability is calculated as the fraction of frames where $\lambda_p$ value is higher than 0.5, using the frames in the last 50 ns of all simulations corresponding to the main cluster of each conformation. The bottom panel shows the surface representations of the protein colored by charge for both conformations and at both pH values. All charged residues are considered for the representation, and histidines are colored by protonation probability (equivalent to positive charge probability).

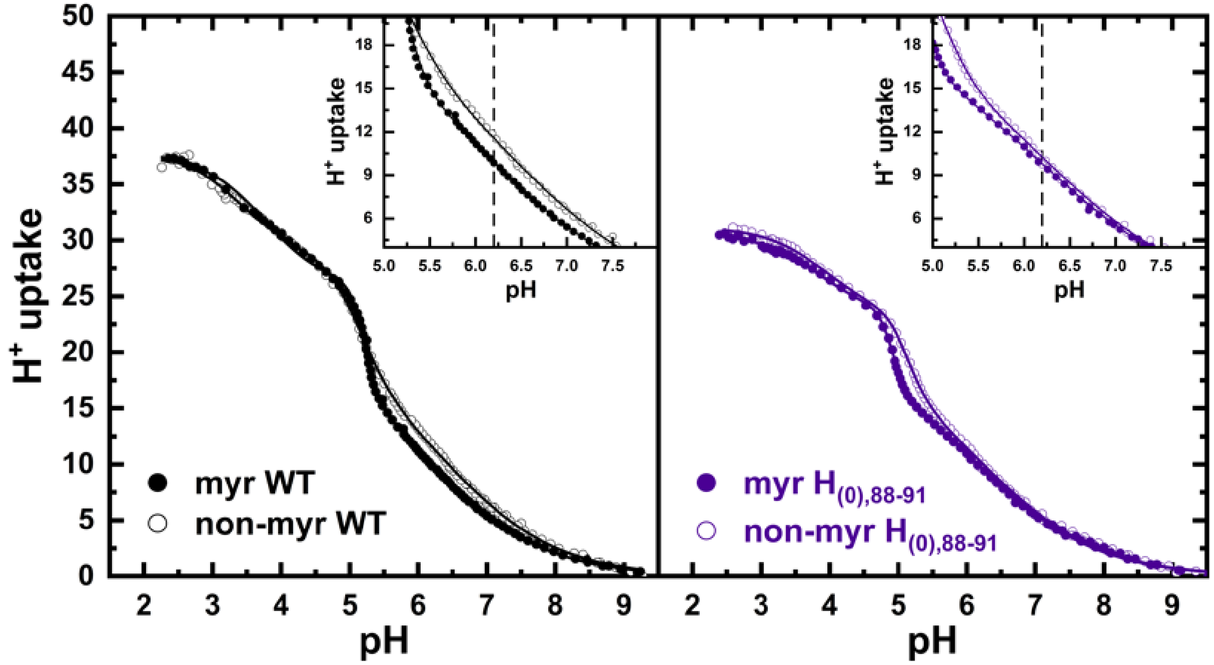

**Figure S6. Experimental proton uptake by wild type and $H_{(0),88\text{-}91}$ mutant of hisactophilin.** Plots are obtained by HCl titration of myristoylated (filled symbols) and non-myristoylated (open symbols) variants of wild type (left) and $H_{(0),88\text{-}91}$ mutant (right). Insets depict a zoom over the pH range of interest, and the 6.2 value is signalled by a dashed line.

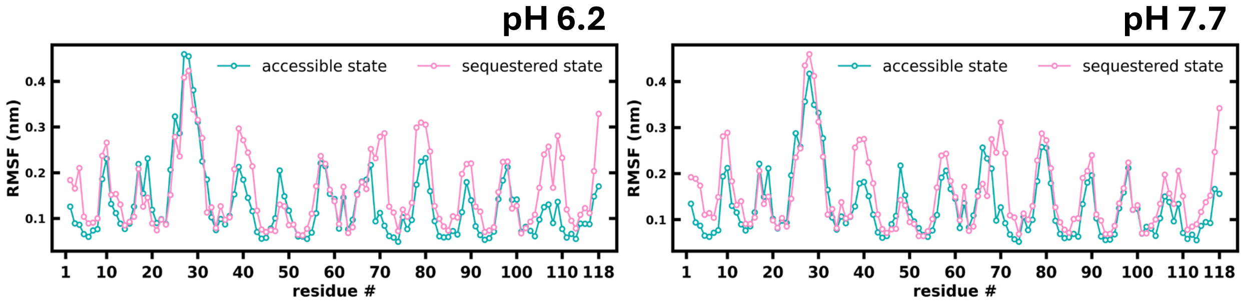

**Figure S7. Root median square fluctuation obtained in the constant pH molecular dynamics simulations.** For the RMSF calculations the residues are represented by the geometric center of their heavy atoms to effectively captures both backbone and side chain dynamics. Values are obtained for the last 50 ns of simulations at pH 6.2 (left panel) and 7.7 (right panel), combining all the frames corresponding to the main cluster of each conformation.

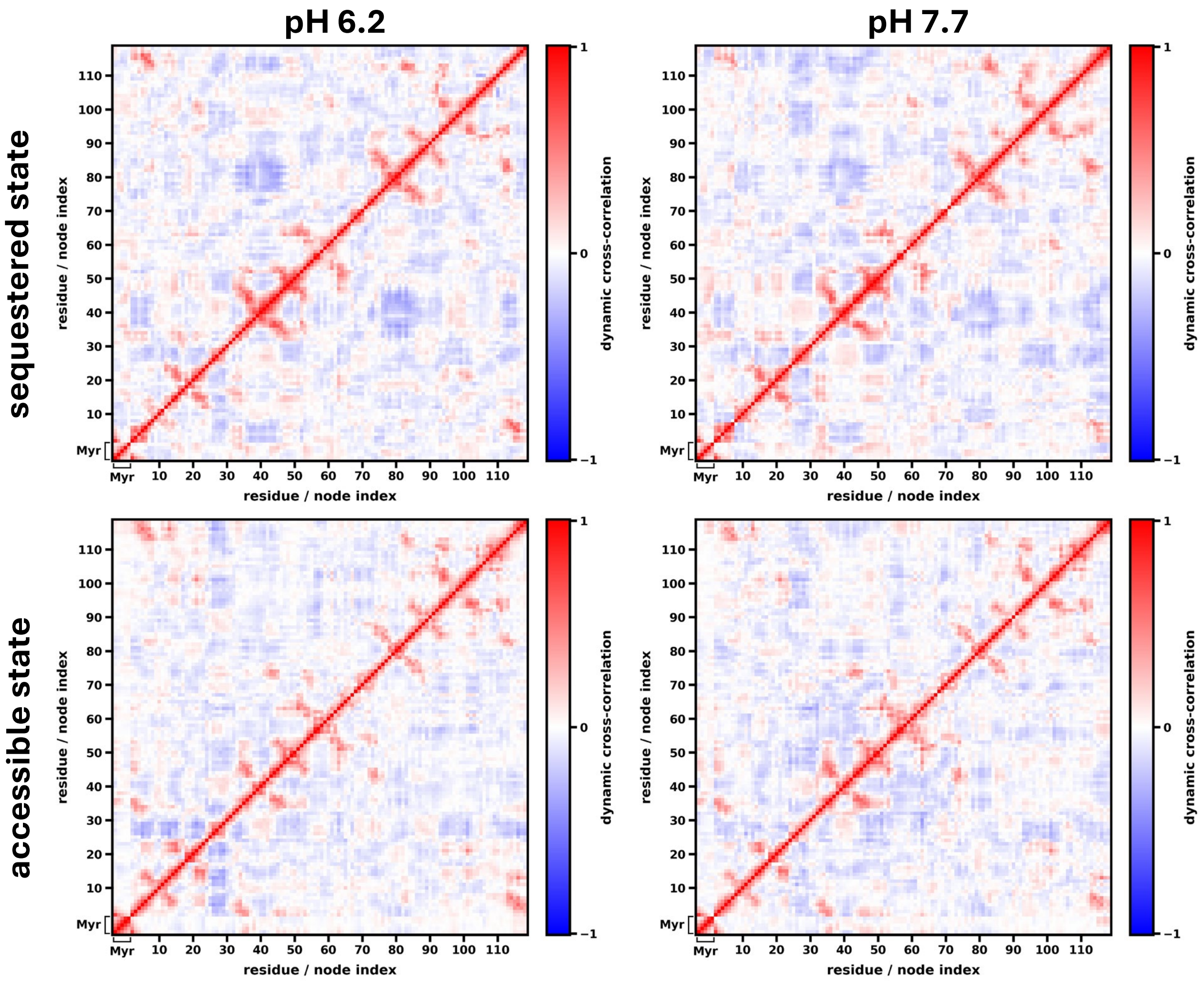

**Figure S8. Dynamic cross-correlation obtained in the constant pH molecular dynamics simulations.** For the cross-correlation calculations the residues are represented by the geometric center of their heavy atoms to effectively captures both backbone and side chain dynamics. Values are obtained for the last 50 ns of simulations at pH 6.2 (left panel) and 7.7 (right panel), combining all the frames corresponding to the main cluster of each conformation.

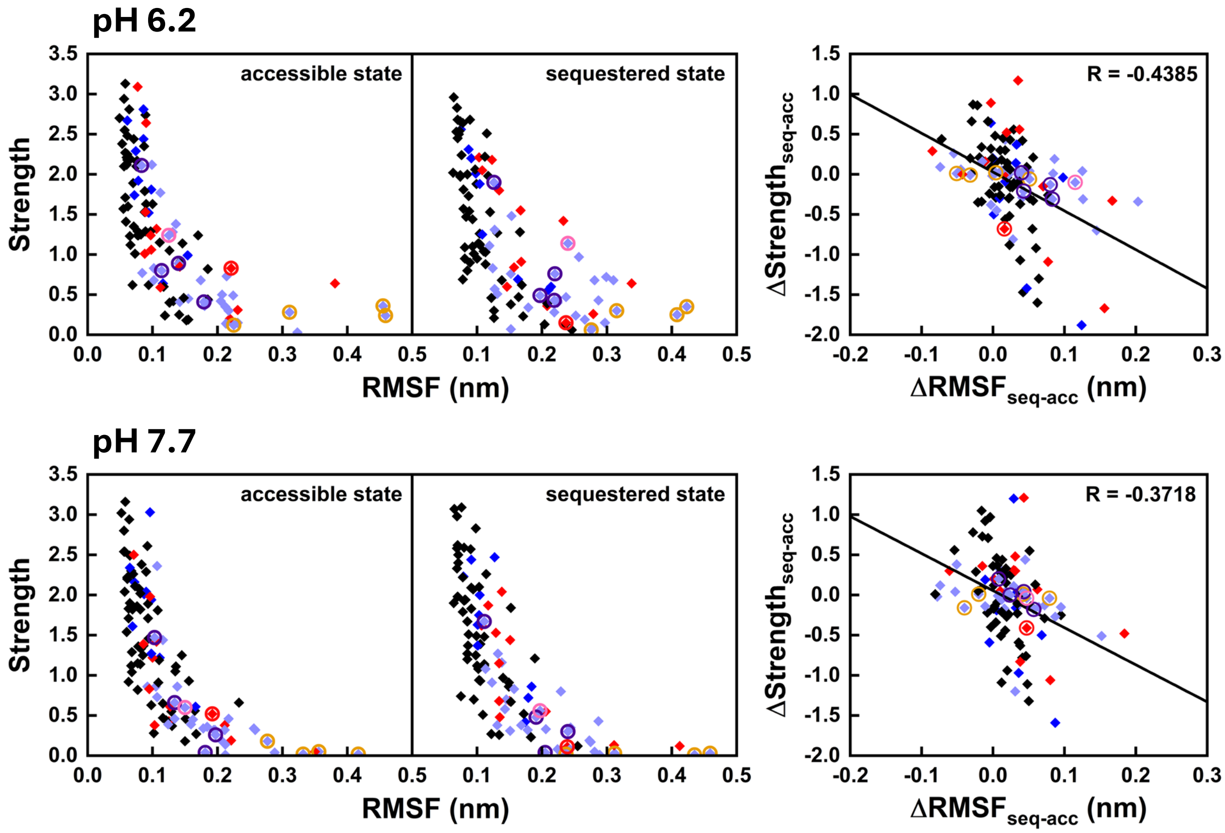

**Figure S9. Inverse correlation between RMSF and Strength.** For the parameters calculations the residues are represented by the geometric center of their heavy atoms to effectively captures both backbone and side chain dynamics. Strength values correspond to the sum of a given node's edge weights (|DCC|). For the RMSF calculation, the last 50 ns of all simulations at pH 6.2 (top panels) and 7.7 (bottom panels) were used, combining all the frames corresponding to the main cluster of each conformation. For the Strength calculation, all replicates were separated into five independent groups with an approximately equal number of frames, with each group using frames from the last 50 ns of simulations corresponding to the main cluster of each conformation. Symbol are colored based on their charge as red (negative charge), blue (positive charge), black (neutral), while histidines are highlighted in light blue. Circles surrounding symbols depict residues mutated in this work (27–31 in orange, 88–91 in purple, 106 in pink, and 57 in red). The Pearson correlation coefficient of the differences between states is included in its corresponding panel.

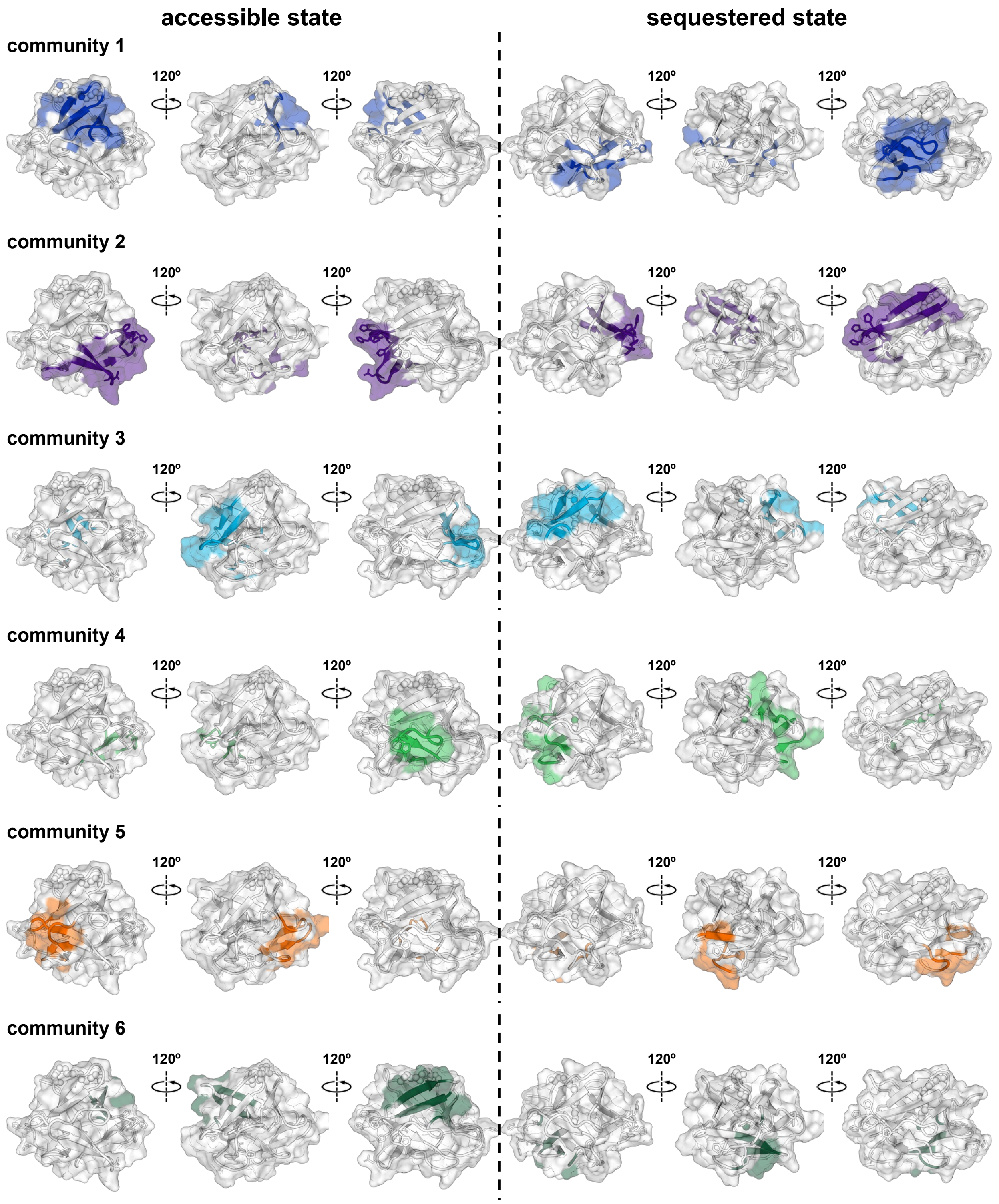

**Figure S10. Residue communities detected from the constant pH molecular dynamics simulations at pH 6.2.** Each panel depicts hisactophilin's structure in each conformational state with individual communities of nodes colored. Communities of nodes were identified using a consensus approach with the Leiden algorithm and are sorted from largest to smaller, based on the number of nodes. Community analysis was performed on the last 50 ns of all simulations corresponding to the main cluster of each conformation.

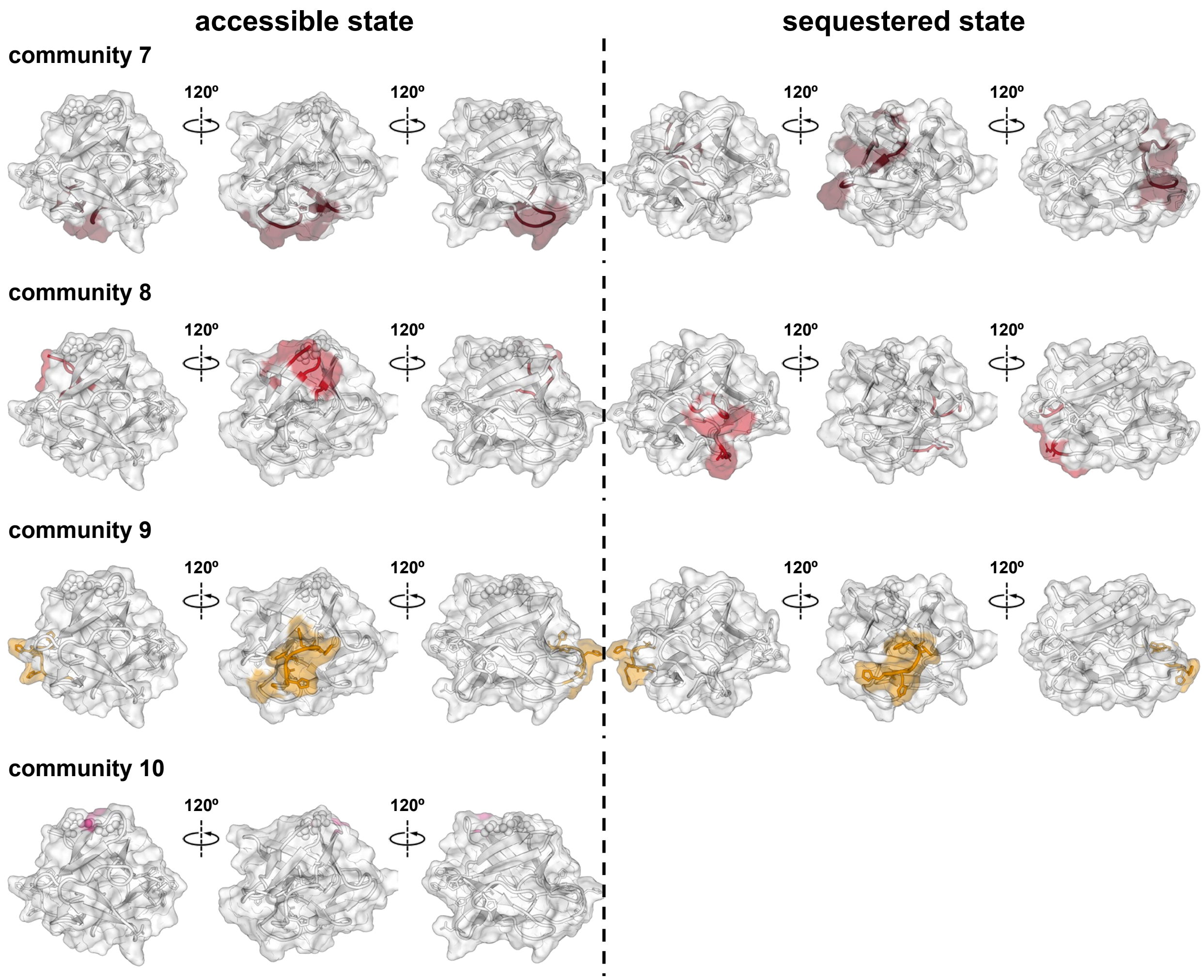

**Figure S10 (cont.).**

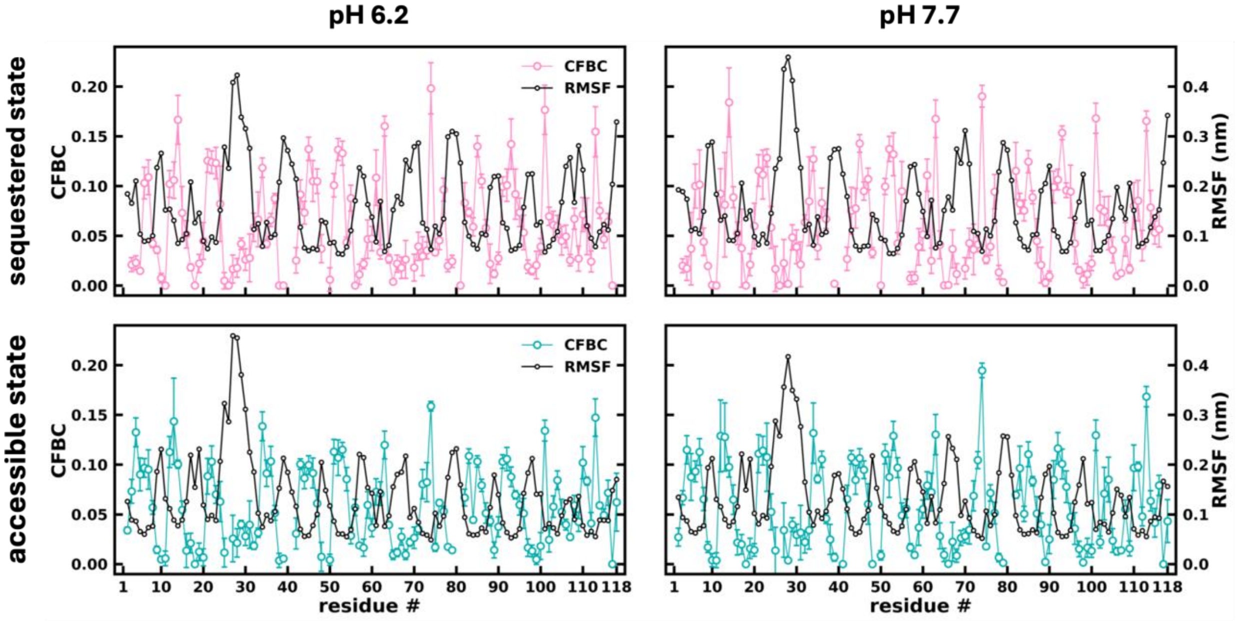

**Figure S11. CFBC increases in regions of low RMSF.** Comparing CFBC and RMSF, both represented by the geometric center of their heavy atoms to effectively captures both backbone and side chain dynamics. RMSF values are obtained for the last 50 ns of simulations at pH 6.2 (left panel) and 7.7 (right panel), combining all the frames corresponding to the main cluster of each conformation. CFBC values are calculated by separating all replicates into five independent groups with an approximately equal number of frames, with each group using frames from the last 50 ns of simulations corresponding to the main cluster of each conformation.
